# Active site closure stabilizes the backtracked state of RNA polymerase

**DOI:** 10.1101/410746

**Authors:** Matti Turtola, Janne J. Mäkinen, Georgiy A. Belogurov

**Author notes:** To whom correspondence should be addressed. University of Turku, Dept. of Biochemistry, FIN-20014, Turku, Finland.

## Abstract

All cellular RNA polymerases (RNAP) occasionally backtrack along the template DNA as part of transcriptional proofreading and regulation. Here, we studied the mechanism of RNAP backtracking by one nucleotide using two complementary approaches that allowed us to precisely measure the occupancy and lifetime of the backtracked state. Our data show that the stability of the backtracked state is critically dependent on the closure of the RNAP active site by a mobile domain, the trigger loop (TL). The lifetime and occupancy of the backtracked state measurably decreased by substitutions of the TL residues that interact with the nucleoside triphosphate (NTP) substrate, whereas amino acid substitutions that stabilized the closed active site increased the lifetime and occupancy. These results suggest that the same conformer of the TL closes the active site during catalysis of nucleotide incorporation into the nascent RNA and backtracking by one nucleotide. In support of this hypothesis, we construct a model of the 1-nt backtracked complex with the closed active site and the backtracked nucleotide in the entry pore area known as the E-site. We further propose that 1-nt backtracking mimics the reversal of the NTP substrate loading into the RNAP active site during on-pathway elongation.

## INTRODUCTION

RNA polymerase (RNAP) mediates the synthesis of an RNA copy of the template DNA—the first and often decisive step in gene expression. All RNAPs transcribing cellular genomes are multisubunit enzymes that share homologous catalytic cores. The bacterial RNAP, a five-subunit complex, ααββ’ω, is the simplest model system used for studying the fundamental mechanistic properties of all multisubunit RNAPs.

The cycle of nucleotide incorporation by RNAP involves binding of the substrate NTP into the active site followed by the closure of the active site by a mobile domain called the trigger loop (TL)(Vassylyev *et al*, 2007b; Wang *et al*, 2006). In the closed conformation, the TL forms a triple helical bundle with another important active site element, the bridge helix (BH)(Gnatt *et al*, 2001; Hein & Landick, 2010). Closure of the active site accelerates catalysis 10,000 fold by positioning the triphosphate moiety of the substrate NTP inline for an attack by the 3’OH group of the RNA. Incorporation of the NMP into the nascent RNA results in a pre-translocated complex where the newly synthesized RNA 3’ end occupies the substrate site (aka the A site). The subsequent post-catalytic relaxation into the catalytically competent post-translocated state involves the release of pyrophosphate, opening of the active site, and forward translocation along the template DNA, which frees the A site for binding of the next NTP.

RNAP translocation along the DNA is a Brownian process (Yager & von Hippel, 1991; Guajardo & Sousa, 1997; Bar-Nahum *et al*, 2005; Abbondanzieri *et al*, 2005; Bai *et al*, 2004), thus the pre-translocated RNAP may move backward (backtrack) instead of translocating forward. Backtracking by one nucleotide necessitates the 3´ end NMP of the nascent RNA to unpair from the template DNA in the A site (Komissarova & Kashlev, 1997b, 1997a; Nudler *et al*, 1997) and to relocate elsewhere in the active site (Wang *et al*, 2009; Sekine *et al*, 2015). Backtracking by many nucleotides extrudes the unpaired nucleotides into the secondary channel of RNAP (Cheung & Cramer, 2011), a route for NTP substrate entry into the active site. The probability to backtrack is typically low but may be significant (10-20%) in certain sequence contexts (Artsimovitch & Landick, 2000). In addition, the probability of backtracking is increased if a roadblock, such as a DNA bound protein (Epshtein *et al*, 2003; Dangkulwanich *et al*, 2013; Kotlajich *et al*, 2015) or a lesion (Charlet-Berguerand *et al*, 2006; Xu *et al*, 2017), impedes forward movement. Misincorporation events also accelerate backtracking by easing the unpairing of the RNA 3’ end, which is a pre-requisite for backtracking (Da *et al*, 2016).

Backtracking often leads to pauses that last from seconds to minutes (Shaevitz *et al*, 2003; Abbondanzieri *et al*, 2005; Galburt *et al*, 2007) and therefore crucially influences gene regulation by modulating the rate of transcription (Adelman *et al*, 2005; Proshkin *et al*, 2010; Perdue & Roberts, 2010; Jonkers & Lis, 2015; Artsimovitch & Landick, 2000). Backtracking is also an essential component of some transcription-coupled DNA repair pathways (reviewed in (Belogurov & Artsimovitch, 2015)) and transcription proofreading (Erie *et al*, 1993; Zenkin *et al*, 2006; Sydow & Cramer, 2009; Imashimizu *et al*, 2015; Bubunenko *et al*, 2017). At the same time, extensive backtracking negatively impacts cell fitness, as it leads to R-loops and double-strand breaks in the DNA upon collisions between RNAP and the replisome (Dutta *et al*, 2011; Sankar *et al*, 2016).

RNAP backtracked by 1 nt can recover into a catalytically-competent post-translocated state by two successive rounds of forward translocation or by endonucleolytically cleaving two nucleotides from the 3’ end of the nascent RNA. The latter reaction is catalyzed by the RNAP active site (Orlova *et al*, 1995) with the assistance of the dissociable factors GreA and B in bacteria (Borukhov *et al*, 1993; Laptenko *et al*, 2003), TFS in archaea (Hausner *et al*, 2000; Fouqueau *et al*, 2017) and eukaryotes (Izban & Luse, 1992; Reines, 1992), or the mobile domains of specialized subunits in the case of the eukaryotic Pol I and Pol III (Ruan *et al*, 2011). The predominant recovery mechanism likely differs depending on the species, transcribed sequence, and the availability of the cleavage stimulating factors. TECs backtracked by many nucleotides are less likely to recover by thermally driven forward translocation (Lisica *et al*, 2016) and thus presumably rely on the endonucleolytic cleavage of the nascent RNA or forward biasing molecular motors such as trailing RNAPs (Epshtein *et al*, 2003) and ribosomes (Proshkin *et al*, 2010) or the Mfd translocase (Park *et al*, 2002) for recovery. RNAP backtracked by 1 nt arguably represents the mechanistically simplest, physiologically most prevalent, and functionally most important off-pathway state that does not involve the rearrangement of RNAP domains relative to the on-pathway post-and pre-translocated states (Sekine *et al*, 2015). Here, we employed a transcription elongation complex (TEC), where a fluorescent nucleobase 2-aminopurine (2AP) at the 3’ end of the nascent RNA forms an unusual Watson-Crick base pair with the thymine in the template DNA, to induce backtracking and to study its mechanism. Our data suggest that closure of the active site by TL strongly stabilizes the backtracked state and support a plausible model of the backtracked state that accounts for the experimental data but is distinct from the backtracked state observed in crystals (Wang *et al*, 2009; Sekine *et al*, 2015). We further propose a mechanism in which 1-nt backtracking mimics the reversal of the NTP substrate loading into the RNAP active site during on-pathway elongation.

## RESULTS

### TEC with a 2AP at the 3’ end of the nascent RNA is fractionally inactivated in a Mg^2+^ dependent manner

To study the properties of the backtracked state, we constructed a backtracked prone TEC using an RNA primer with a 2-aminopurine (2AP) in its 3’end. 2AP is an adenine analogue, which forms an unusual Watson-Crick base pair with thymine in the template DNA strand (**Figure 1A,B**), thereby increasing the propensity of RNAP to backtrack (see below). In addition, 2AP displays low fluorescence when stacked with neighboring bases and high fluorescence when unstacked (Hardman & Thompson, 2006), allowing us to monitor the separation of the RNA 3’ end from the template DNA. The TEC (2AP-TEC) was assembled on a synthetic nucleic acid scaffold and contained the fully complementary transcription bubble flanked by 22 and 18 nucleotide DNA duplexes upstream and downstream, respectively. The annealing region of a 17-nucleotide RNA primer was 11 nucleotides, permitting the TEC to adopt the post-translocated, pre-translocated and 1-nt backtracked states. The RNA primer was 5’ labeled with the infrared fluorophore ATTO680 to monitor RNA extension by denaturing PAGE.

**Figure 1.**
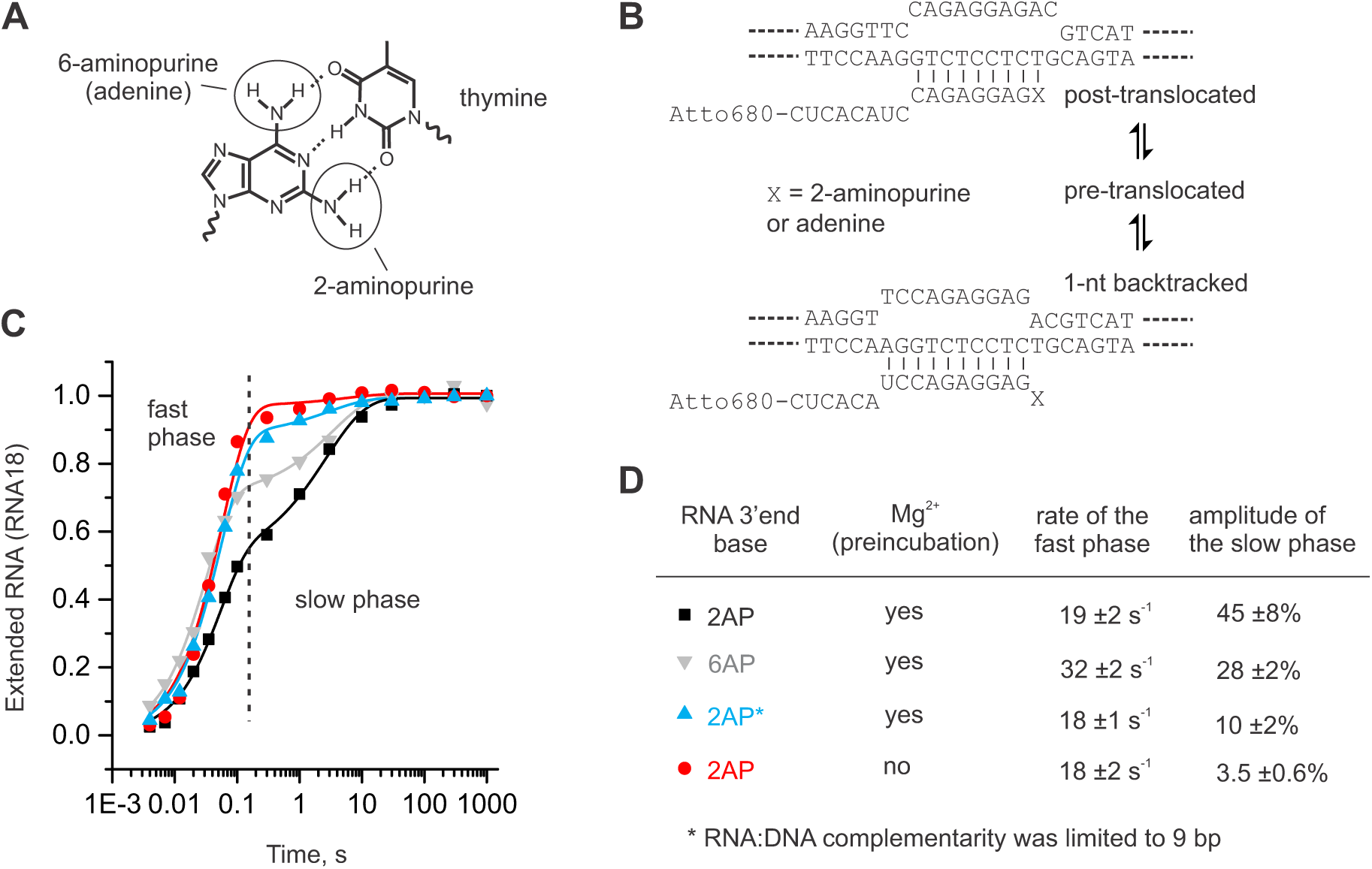
TEC with a 2-aminopurine at the 3’ end of the nascent RNA is fractionally inactivated in a Mg^2+^ dependent manner. **(A)** 2-aminopurine (2AP) forms an unusual Watson-Crick base pair with thymine. **(B)** The schematics of the post-translocated and backtracked states of the TECs used in the experiments in (C), interconversion between these states likely takes place via the pre-translocated state as is shown in the scheme. **(C)** Kinetics of CMP incorporation. TECs contained 2AP (black, blue and red) or adenine (grey) as the base at the 3’ end of the RNA. TECs were either preincubated with 1 mM Mg^2+^ and supplemented with an equal volume of 400 µM CTP containing 1 mM Mg^2+^ (black, blue, grey) or a Mg^2+^-free TEC was supplemented with an equal volume of 400 µM CTP containing 2 mM Mg^2+^ (red). The final reaction conditions (200 µM CTP, 1 mM Mg) were identical in all experiments. The RNA:DNA complementarity was 11 bp (black, grey, red) or 9 bp (blue). The oligonucleotides used for assembling the TECs are shown in **Supplementary Figure S1**. The data points are averages of 2-3 independent experiments (error bars are omitted for clarity) and the solid lines are the best-fits to a sum of exponential (corresponds to the fast phase) and stretched exponential (corresponds to the slow phase) functions (see Methods for details). **(D)** The rates of the fast phase and the amplitudes of the slow phase inferred from the data in (C).

To determine if the 2AP-TEC is fractionally inactivated, we measured the kinetics of the incorporation of the next nucleotide (CMP) in a rapid chemical quench flow device. The RNA cleavage activity of the 2AP-TEC was very low (see the dedicated section below) permitting the accurate analysis of the RNA extension in the 2AP-TEC supplemented with Mg^2+^ prior to loading into the quench flow instrument. CMP incorporation followed biphasic kinetics with fast (∼20 s^-1^) and slow (∼0.4 s^-1^) phases of approximately equal amplitudes when both TEC and CTP solutions contained 1 mM Mg^2+^ (**Figure 1C**). However, when the reaction was initiated by mixing a Mg^2+^-free solution of 2AP-TEC with a CTP solution containing 2 mM Mg^2+^, the amount of the slow fraction was drastically reduced. The rate of CMP incorporation in the fast fraction matched that of the 2AP-TEC locked in the post-translocated state by limited RNA:DNA complementarity, whereas the rate of CMP incorporation in the slow phase was similar to the recovery rates of the short sequence-specific pauses observed in the single-molecule experiments (Herbert *et al*, 2006; Shundrovsky *et al*, 2004). We concluded that (*i*) the 2AP-TEC is fractionally and reversibly inactivated, (*ii*) the inactivation is possibly due to the formation of the backtracked state, and (*iii*) the formation of the inactivated state requires pre-incubation with Mg^2+^.

Importantly, using an RNA primer with an adenine (6AP) at the RNA 3’ end also resulted in a TEC with biphasic CMP incorporation kinetics but the fraction of the slow phase decreased ∼1.5 fold comparing with the 2AP-TEC (**Figure 1C,D; Supplementary Table S1**). The slow fraction of 6AP-TEC recovered with the same or slower rate (median recovery time 2.7 ±0.6 s) than that of the 2AP-TEC (median recovery time 1.9 ±0.3 s) suggesting that the 2AP increases the fraction of the inactivated state by increasing the entry rate and without significantly altering the stability of the inactivated state (i.e. the recovery rate). The increased backtracking rate in the 2AP-TEC is not unexpected considering that (*i*) the 2AP-T base pair is weaker than 6AP-T base pair despite featuring the same number of hydrogen bonds (Eritja *et al*, 1986) and (*ii*) the proteinaceous environment of the RNAP active site possibly amplifies the differences in the base pair stabilities as a part of its general capacity to facilitate the discrimination between the NTP substrates. Overall, we concluded that the 2AP-TEC is a good model system for studying backtracking because it qualitatively matches the behavior of the fully native system but additionally allows monitoring of the unstacking of the RNA 3’ end by measuring the fluorescence.

### Formation of the inactivated state increases the fluorescence of the 2-aminopurine at the 3’ end of the nascent RNA

The 2AP-TEC assembled in the absence of Mg^2+^ displayed low fluorescence, but the addition of 1 mM Mg^2+^ increased the fluorescence 6-fold, suggesting that a fraction of the 2AP moieties becomes unpaired and separated from the penultimate guanine base in the latter case (**Figure 2A**). The increase in the fluorescence upon addition of Mg^2+^ followed a single exponential function with a rate constant of ∼2 s^-1^ (**Figure 2B**). The rate of the fluorescence increase was arguably too slow to describe any step in the on-pathway transcript elongation or Mg^2+^ binding, and therefore likely reflected the formation of an off-pathway state. The increase in the fluorescence was coupled to an increase in fluorescence anisotropy (**Figure 2B**), suggesting that a fraction of the 2AP moieties also became more ordered. The extension of the TEC with CMP reduced the fluorescence to the Mg^2+^-free level **(Figure 2A)**, presumably due to quenching of the 2AP fluorescence by the neighboring guanine and cytosine bases within the RNA:DNA hybrid. The decrease in TEC fluorescence upon CMP incorporation occurred with a similar rate to that of the slow phase of CMP incorporation in rapid chemical quench flow experiments (**Figure 2C**). These results suggest that the high fluorescent state corresponds to the inactivated state that forms reversibly and is stabilized by a Mg^2+^ ion. Interestingly, the endonucleolytic cleavage of the Mg^2+^-supplemented 2AP-TEC by GreA decreased the fluorescence more than twofold demonstrating that the 2AP fluorescence in the pGp2AP dinucleotide is lower than in the inactivated state (**Supplementary Figure S2**). The latter observation lends credence to the hypothesis that the 2AP is forced to unstuck from the penultimate guanine base in the inactivated state.

**Figure 2.**
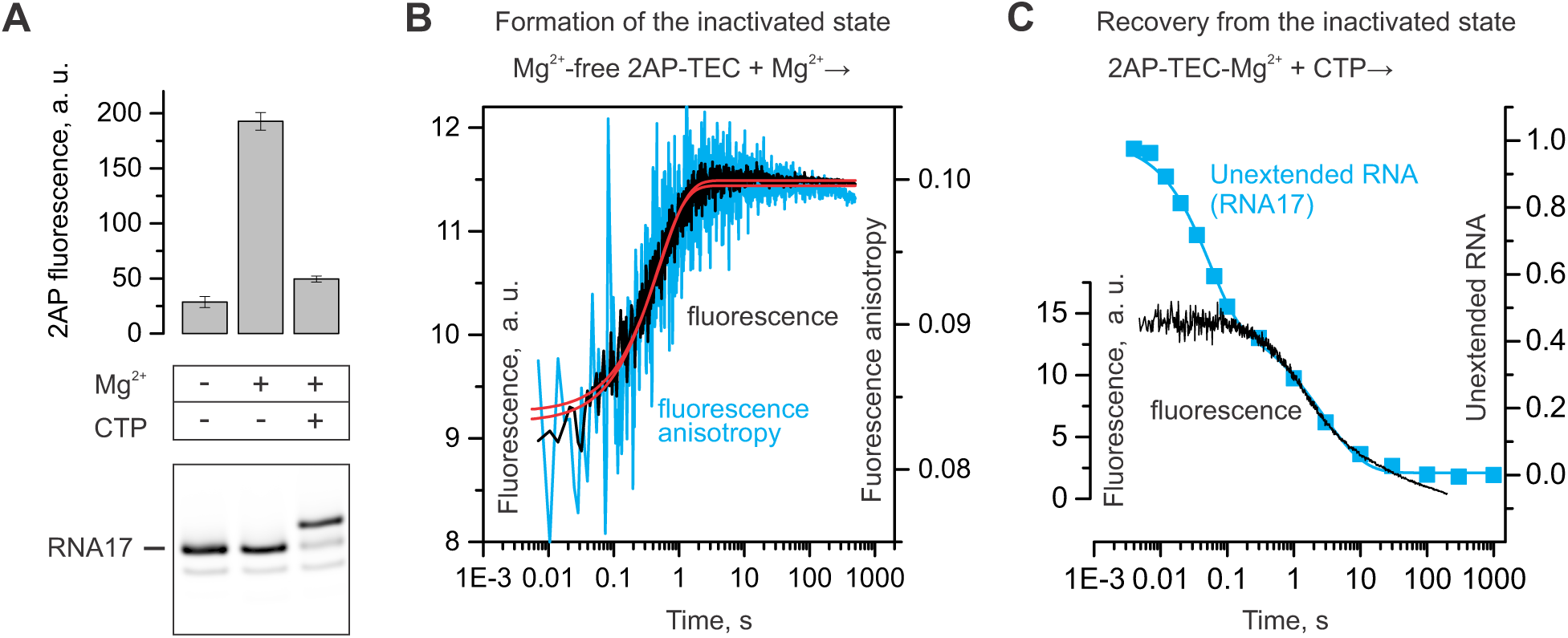
Formation of the inactivated state increases the fluorescence of the 2-aminopurine at the 3’ end of the nascent RNA. A schematic of the 2AP-TEC used in these experiments is shown in **Figure 1B**. **(A)** Addition of Mg^2+^ to a Mg-free 2AP-TEC increases the fluorescence, further addition of CTP decreases the fluorescence to the near initial level as recorded at equilibrium conditions using a fluorometer. Error bars show the range of duplicate measurements. A denaturing PAGE gel of the RNA in the 2AP-TECs is shown below the bar graph. **(B)** Time resolved measurements of the increase in fluorescence upon the addition of Mg^2+^ to a Mg^2+^-free 2AP-TEC. The increase in fluorescence (black trace) and fluorescence anisotropy (blue trace) are well described by a single exponential function (red best-fit lines) with the rate constant 2.0 ±0.03 and 2.2 ±0.1 s^-1^, respectively. **(C)** Time resolved measurements of the decrease in fluorescence (black trace) upon the addition of CTP to the 2AP-TEC pre-incubated with 1 mM Mg^2+^. Fluorescence decreases with the median time of 3.8 ±0.05 s as inferred by fitting to a stretched exponential function (see Methods). The decrease in the fraction of the unextended RNA is shown for comparison (blue squares). The decrease in the fraction of RNA17 corresponds to the increase in the fraction of RNA18 presented in **Figure 1C** (black squares). The slow fraction of RNA17 reacts with CTP with the median time of 3.8 ±0.1 s.

### TEC inactivation corresponds to backtracking by one nucleotide

Next, we mapped the translocation register of the inactivated state of the 2AP-TEC by disrupting the base pairing within the RNA:DNA hybrid and the downstream DNA. Backward translocation by one or more registers involves reannealing of the RNA with the template DNA (tDNA) at the upstream edge of the RNA:DNA hybrid. Shortening the RNA:DNA complementarity selectively destabilizes the backtracked states and biases the TEC towards the pre-and post-translocated states. Shortening the RNA:DNA from 11 to 10 bp reduced the amplitude of the Mg^2+^ induced increase in the 2AP fluorescence more than twofold (U to A substitution, **Figure 3A**), whereas further shortening to 9 bp practically eliminated this effect (UC to AA substitutions, **Figure 3A**).

**Figure 3.**
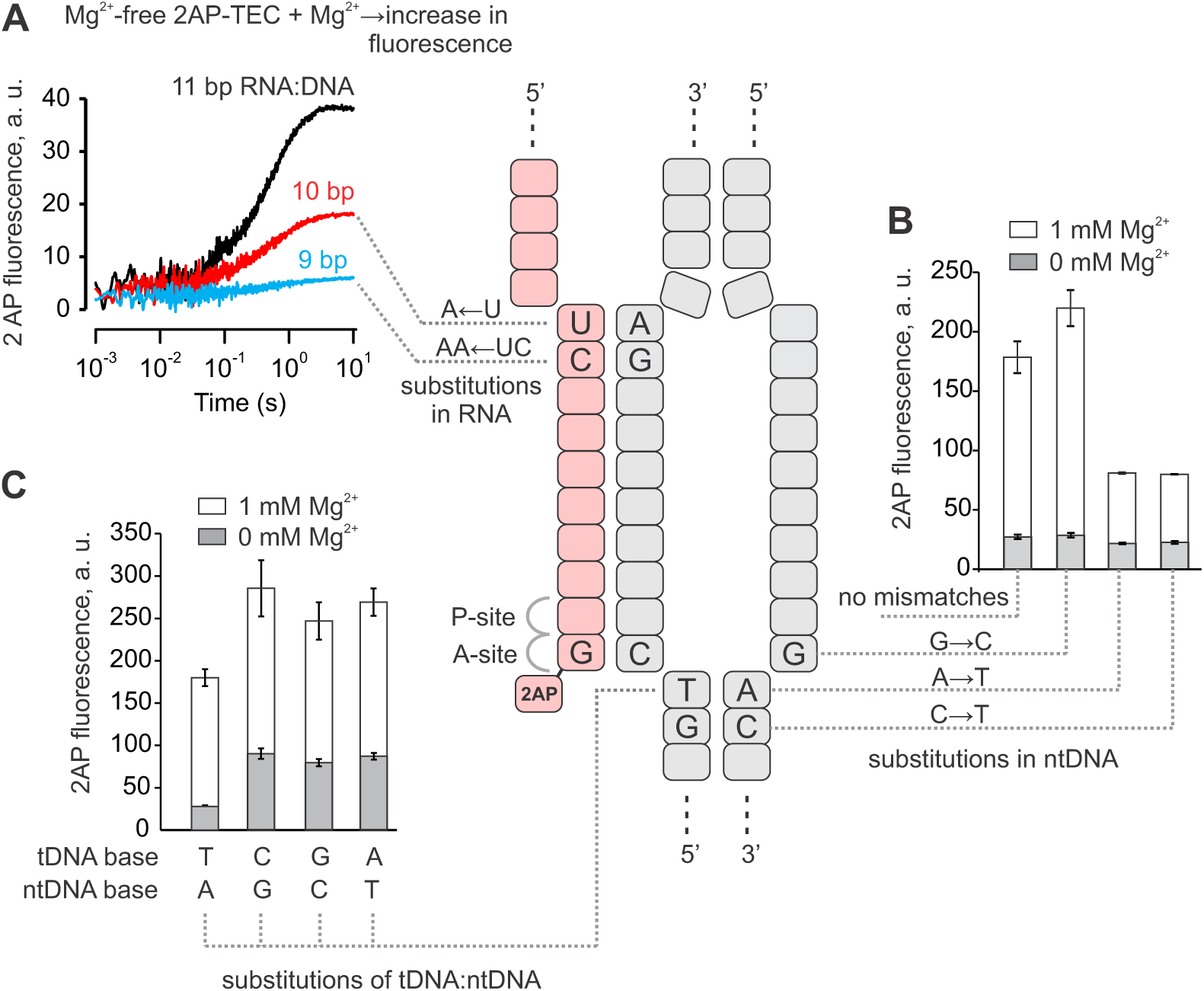
Mapping the translocation register of the inactivated state in 2AP-TEC. The schematic in the center depicts the 2AP-TEC backtracked by one nucleotide. **(A)** The effects of the RNA:DNA mismatches at the upstream edge of the transcription bubble on the amplitude of the increase in fluorescence upon mixing of the Mg^2+^-free 2AP-TEC with an equal volume of a 2 mM Mg^2+^ solution in a stopped flow instrument. **(B)** The effects of tDNA:ntDNA mismatches at the downstream edge of the transcription bubble on the fluorescence of the 2AP-TEC in the presence and absence of 1 mM Mg^2+^. **(C)** The effects of the 2AP-tDNA mismatches on the fluorescence of the 2AP-TEC in the presence and absence of 1 mM Mg^2+^. Error bars show the range of duplicate measurements. The oligonucleotides used for assembling the TECs are shown in **Supplementary Figure S1**.

Next, backtracking by one or more registers involves reannealing of the downstream DNA. Eliminating the DNA:DNA complementarity at the position corresponding to the RNA 3’ NMP (A to T substitution, **Figure 3B)** or immediately downstream of the RNA 3’ end (C to T substitution, **Figure 3B)** reduced the 2AP-TEC fluorescence by destabilizing the backtracked state. In contrast, eliminating the DNA:DNA complementarity immediately upstream of the RNA 3’ NMP (G to C substitution, **Figure 3B)** marginally increased the 2AP-TEC fluorescence by selectively destabilizing the long-backtracked states (≥ 2 nt). We suggest that the 2AP fluorescence is lower in the long-backtracked states (≥ 2 nt) because 2AP is not forced to unstack from the penultimate guanine, consistently with the observation that the 2AP fluorescence in the pGp2AP nucleotide is lower than in the 1-nt backtracked state (**Supplementary Figure S2**). Finally, backtracking involves the separation of the RNA 3’ end from the template DNA. Improving the interaction of the RNA 3’ end with the template DNA by replacing the 2AP with adenine diminished the fraction of the inactivated state (**Figure 1C,D**). In contrast, compromising the interactions of the RNA 3’ end with the template DNA by replacing a thymine with a cytosine, adenine or guanine increased the fraction of the inactivated state, as judged from the 40-60% higher total fluorescence of the 3’ mismatched TECs (**Figure 3C**). Overall, the analysis presented above strongly suggests that the high-fluorescence, Mg^2+^ dependent inactivated state formed in the 2AP-TEC is a 1-nt backtracked state.

### Active site closure by the trigger loop stabilizes the 1-nt backtracked state

We have previously demonstrated that resting TECs are predominantly post-translocated and that the formation of a measurable fraction of a pre-translocated TEC requires the closure of the active site by the TL folding into the TH (Malinen *et al*, 2012). Here, we investigated the effects of perturbations affecting the TH-TL equilibrium on the formation of the backtracked state. The Mg^2+^-dependent increase in the fluorescence was largely abolished in a 2AP-TEC assembled with an RNAP lacking the entire TL (δTL RNAP) (**Figure 4A**), suggesting that the TL is a necessary component of the Mg^2+^-stabilized backtracked state. Similarly, the addition of 10 µM of the antibiotic streptolydigin (Stl), which traps the TL in the open conformation (Temiakov *et al*, 2005; Vassylyev *et al*, 2007b), reduced the fluorescence to the level of the δTL 2AP-TEC (**Figure 4A;** two rightmost bars). The result is consistent with earlier observations that Stl disfavors the backtracked state (Tuske *et al*, 2005). We used the Stl-hypersensitive β´N792D RNAP (Temiakov *et al*, 2005; Malinen *et al*, 2014) in the latter experiment because the high concentration of Stl (500 µM) required for the inhibition of the wild-type RNAP interfered with the fluorescence measurements. The β’N792D substitution improves Stl binding to *Escherichia coli* (*Eco*) RNAP (Temiakov *et al*, 2005) but does not affect the nucleotide addition and the translocation rates (Malinen *et al*, 2014).

**Figure 4.**
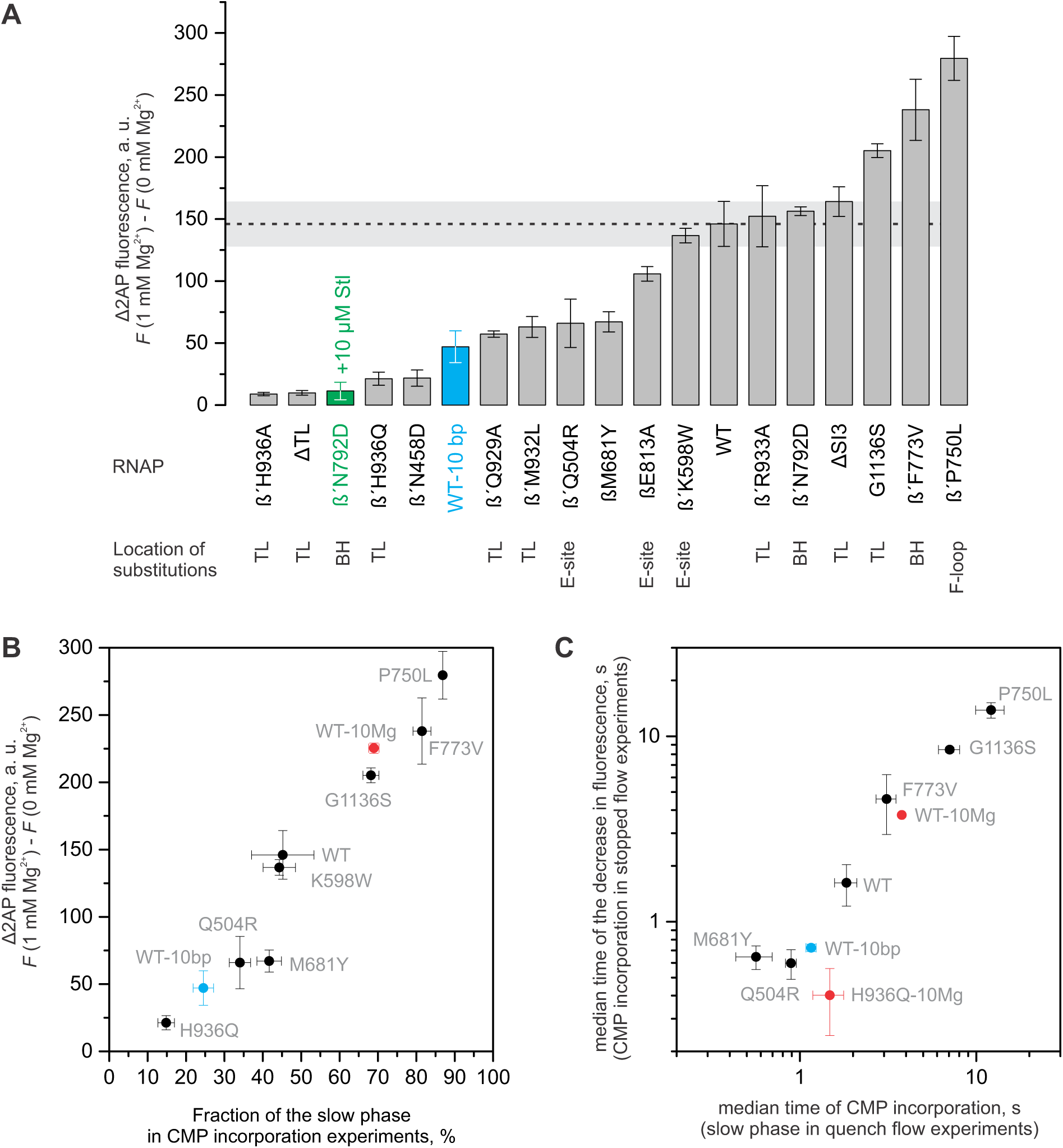
Effects of the amino acid substitutions in the RNAP active site on the occupancy (A, B) and stability (C) of the 1-nt backtracked state. Most of the experiments were performed at 1 mM Mg^2+^ using 2AP-TECs with 11bp of RNA:DNA complementarity. A subset of experiments was performed at 10 mM Mg^2+^ (red), using TECs with reduced RNA:DNA complementarity (blue) or in the presence of streptolydigin (green). Error bars represent the range of duplicate measurements or standard deviations from multiple experiments. **(A)** The increase in fluorescence was recorded upon the addition of 1 mM Mg^2+^ to the 2AP-TECs assembled in the Mg^2+^-free buffer. The observations are arranged in order of increasing amplitude from left to right. The horizontal dashed line depicts the amplitude of the fluorescence increase in the wild-type 2AP-TEC ±SD (horizontal grey bar). **(B)** The correlation of the amplitude of Mg^2+^ induced increase in fluorescence with the fraction of the slow phase in the CMP incorporation experiments. **(C)** The correlation of the recovery kinetics of the backtracked state in the fluorescence and the quench flow experiments.

We next examined the occupancies of the backtracked state in a set of active site variants by estimating the fraction of the slow phase in CMP incorporation experiments and/or by measuring the increase in the fluorescence of a Mg^2+^-free 2AP-TEC upon addition of 1 mM Mg^2+^. The former assay only works well with RNAP variants that have a largely uncompromised nucleotide addition activity that ensures good separation of the fast and the slow phases of CMP incorporation. In contrast, the fluorescence method allowed the estimation of the occupancy of the backtracked state in variant RNAPs with a compromised catalytic activity but may possibly report the direct effects of the substitutions on the 2AP fluorescence in some cases. That said, the two methods reported qualitatively similar effects of the amino acid substitution on the occupancy of the backtracked state in all cases, where the two methods could be applied to the same variant RNAP (**Figure 4B**). Individual substitutions of amino acid residues in RNAP, which are predicted to stabilize (β´P750L, β´F773V, (Malinen *et al*, 2014) and β´G1136S (Mejia *et al*, 2015)) or destabilize (β´Q929A (Basu *et al*, 2014), β´M932L and β´H936A (Vassylyev *et al*, 2007b)) the TH, enhanced or reduced, respectively, the Mg^2+^-dependent increase in the fluorescence (**Figure 4A,B**). For example, a substitution of the β´His^936^, which as part of TH interacts with the ultimate phosphate of the pre-translocated RNA, virtually eliminated the Mg^2+^ dependent increase in the fluorescence. In contrast, a 2AP-TEC assembled using an RNAP with a β´P750L substitution in the F-loop, that allosterically stabilizes the TH (Malinen *et al*, 2014), displayed a nearly twofold higher Mg^2+^ dependent increase in fluorescence than did the wild-type 2AP-TEC. At the same time, the removal of a lineage-specific insertion domain SI3 (Iyer *et al*, 2003; Lane & Darst, 2010) in the TL or a substitution of the β´Arg^933^, which as part of the TH binds to the β-and γ-phosphates of the substrate NTP (Vassylyev *et al*, 2007b), did not have measurable effects on the 2AP-TEC fluorescence (**Figure 4A**).

It is important to note that the occupancy of the backtracked state is a poor measure of the stability of the backtracked state. The interconversion between the post-and the backtracked states involves equilibria between the backtracked, pre-and post-translocated states. Accordingly, substitutions can potentially affect the occupancy of the backtracked state by solely biasing the equilibrium between the pre-and post-translocated states that is known to be controlled by the opening and closing of the active site (Malinen *et al*, 2012; Feig & Burton, 2010; Larson *et al*, 2012; Malinen *et al*, 2014). At the same time, the recovery rate from the backtracked state is a direct measure of its stability. To assess the effects of amino acid substitutions on the stability of the backtracked state, we determined the recovery rate from the backtracked state by analyzing the CMP incorporation kinetics using quench flow and stopped flow setups. The CMP incorporation assay performed in a quench flow setup natively reports if the nucleotide addition rates by the post-translocated state (fast phase) are sufficiently faster than the recovery rates, so that they do not interfere with the estimation of the recovery rates. At the same time, the fluorescence assay has the advantage of reading the signal exclusively from the backtracked state.

The 2AP-TECs assembled with the variant RNAPs and pre-incubated with 1 mM Mg^2+^ showed biphasic CMP incorporation kinetics, with the fraction of a slow phase in good correlation with the amplitude of the Mg^2+^-dependent increase in the fluorescence (**Figure 4B**). The rate of the slow phase and the rate of the decrease in the 2AP fluorescence upon the addition of CMP also correlated well among the tested 2AP-TECs (**Figures 4C, Supplementary Table S1**). The combination of the two assays allowed for the accurate estimation of the recovery rates for the variants with the catalytic activity and the occupancy of the backtracked state similar or larger than that of the WT enzyme. The substitutions β’F773V, β’G1136S and β’P750L decreased the recovery rate from the backtracked state 3-4, 5-8 and 8-10 fold, respectively, suggesting that they stabilized the backtracked state, in addition to stabilizing the pre-translocated state (Malinen *et al*, 2014). It was difficult to accurately determine the recovery rate for β´H936Q, which had low occupancy of the backtracked state at 1 mM Mg^2+^, necessitating measurements at 10 mM Mg^2+^, where the occupancy of the backtracked state is higher. The recovery rates of β´H936Q determined by CMP incorporation and fluorescence assays were 2.5-and 8-fold faster, respectively, than those of the wild-type RNAP at 10 mM Mg^2+^ (**Figure 4**, **Supplementary Table S1**). These observations suggest that the β´His^936^ stabilizes the backtracked state, presumably as part of the TH. Finally, we did not attempt to determine the recovery rates for the RNAP variants with a significantly compromised catalytic activity (βE813A, β´Q929A, β´H936A, β´ΔTL).

Overall, our data suggest that the TL stabilizes the backtracked state by the same mechanism it stabilizes the pre-translocated state: the TL folds into a helical bundle with the BH, closes the active site and interacts with the penultimate (in the backtracked state) or the 3’ terminal (in the pre-translocated state) RNA nucleotide.

### The closed active site can accommodate the backtracked nucleotide in the E-site

The BH β’F773V, TL β’G1136S and F-loop β’P750L substitutions are located in distinct structural elements but share the markedly reduced forward translocation rate that we proposed is due to the stabilized TH (Malinen *et al*, 2014). Considering the fact that all three substitutions are located beyond the reach of the backtracked nucleotide (**Figure 5A**), their effects on the recovery rate (see above) strongly suggest that the TH stabilizes the 1-nt backtracked state in the 2AP-TEC. However, in the two available crystal structures of the 1-nt backtracked state (Wang *et al*, 2009; Sekine *et al*, 2015) the TL is folded into a conformation that neither closes the active site nor approaches the RNA 3’ end, whereas the nucleobase of the backtracked nucleotide folds upstream towards the βD-loop (**Figure 5C**). Furthermore, superimposition of the backtracked (Sekine *et al*, 2015) and the substrate analogue-bound structures of the TECs and initially transcribed complexes (ITC) revealed that the nucleobase of the backtracked nucleotide occupies approximately the same volume as the β’His^936^ (*Eco* numbering) of the fully helical TH (Vassylyev *et al*, 2007b; Liu *et al*, 2016) or β’Thr^934^ (*Eco* numbering) of the TH with a partially unwound tip (Basu *et al*, 2014; Maffioli *et al*, 2017). These clashes could not be eliminated by adjustments to the side chains of the amino acid residues in the TH.

**Figure 5.**
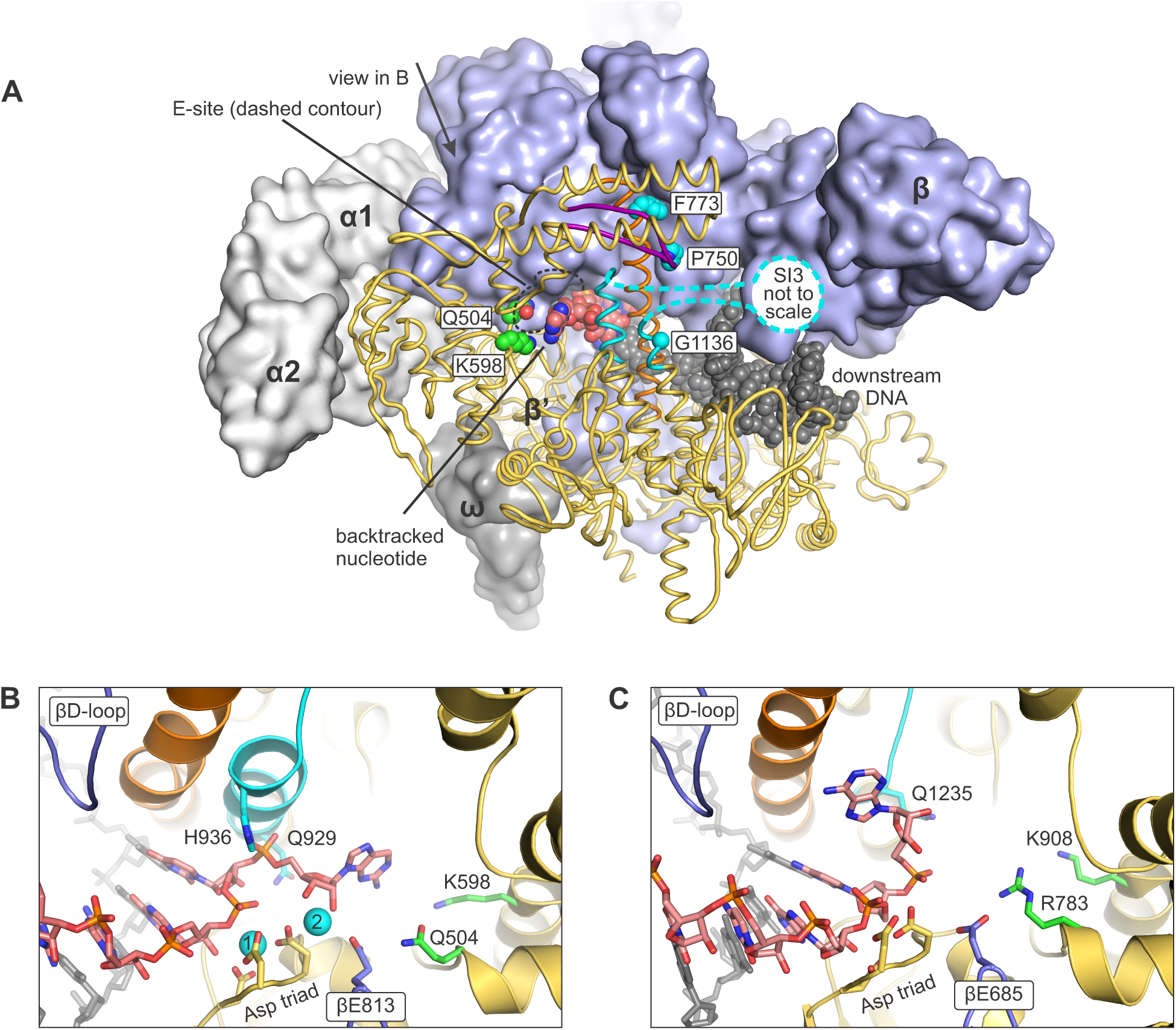
A model of the backtracked state with a closed active site. **(A)** An overview depicting the tentative location of the backtracked nucleotide in the closed active site. The bulk of the β’ is shown as dark yellow cartoons, TL is colored cyan, BH orange, F-loop purple. The nascent RNA (carbons are colored rose), the template DNA (all atoms are colored grey) and the side chains (or α carbon in case of G1136) of the β’ amino acid residues whose substitutions stabilize the closed active site (carbons are colored cyan) or increase the RNA cleavage activity (carbons are colored green) above that expected from the occupancy of the backtracked state are shown as spheres. **(B)** The close up views of the backtracked nucleotide in a model constructed in this study. Only selected regions of β subunit are shown. Mg^2+^ ions are depicted as cyan spheres and numbered according to their relative affinity: higher affinity MG1 and lower affinity MG2. Other colors are as in (A). **(C)** The close up view of the backtracked nucleotide in the crystal structure of *Thermus thermophilus* TEC (PDB ID 4WQS). The homologous residues of *T. thermophilus* are colored as the corresponding residues in the *E. coli* RNAP. No metal ions are observed in the active site of the backtracked *T. thermophilus* RNAP. Figure was prepared using PyMOL Molecular Graphics System, Version 1.8.6.0 Schrodinger, LLC.

The structural and biochemical data can be reconciled in two ways. One possibility is that the active site of the backtracked RNAP in the 2AP-TEC is closed by a TL conformation that is distinct from the TH conformers in the substrate bound TEC and ITC. Another possibility is that the conformation of the RNA 3’ nucleotide is different from that observed in the crystal structure of the backtracked TEC (Sekine *et al*, 2015). We explored the second scenario because the higher than expected RNA cleavage activity of the β´Q504R and β´K598W RNAPs (see below) additionally suggested that the backtracked nucleotide is at least fractionally localized in the area designated as the E-site by Westover et al (Westover *et al*, 2004), far away from the crystallographically observed pose near the βD-loop (**Figure 5**).

Therefore, we performed a manual search for a spatially feasible position of the backtracked nucleotide in the active site closed by the TH. We used the crystal structure of the pre-translocated ITC of *Eco* RNAP (Liu *et al*, 2016) as the initial model because the ITC structures were the only available structures of the *Eco* RNAP with the closed active site at the time of writing. Notably, *Eco* RNAP occasionally backtracks also during the initial transcription (Duchi *et al*, 2016; Lerner *et al*, 2016). The pre-translocated ITC model was *in silico* extended by attaching a 2AP nucleoside monophosphate to the RNA 3’ end, yielding the 1-nt backtracked TEC, and the spatially feasible poses of the backtracked nucleotide were explored by manipulating the torsion angles around the phosphate and C5’ of the backtracked nucleotide and C3’ of the penultimate nucleotide.

Our modeling experiments suggest that the phosphate of the backtracked nucleotide closely approaches the N-terminal arm of the TH approximately midway between the β’Gln^929^ and the β’His^936^ (**Figure 5B**). The ribose occupies the volume between the β’His^936^ and the βGlu^813^, the same area that accommodates pyrophosphate or a pyrophosphate moiety of the NTP (Vassylyev *et al*, 2007b; Malinen *et al*, 2012; Liu *et al*, 2016). The purine nucleobase does not readily fit into the interior of the active site but can be spaciously accommodated (5-10Å from the RNAP sidechains) in the secondary channel area between the β’Arg^731^ and β’Gln^504^ of the active site wall, the β’Arg^933^ and β’Ile^937^ of the TH and β’Lys^598^ at the distant periphery of the active site (**Figure 5B**).

In our model, the nucleobase of the backtracked nucleotide occupies approximately the same volume as the nucleobase of the non-complementary NTP bound in the E-site (Wang *et al*, 2006; Westover *et al*, 2004) and overlaps with the binding site of tagetitoxin (Vassylyev *et al*, 2005). Similarly to tagetitoxin, the hydroxyl groups of the backtracked nucleotide can potentially coordinate an additional Mg^2+^ ion (MG2 in **Figure 5B**), together with the βGlu^813^. Binding of an additional Mg^2+^ ion can possibly explain the dependence on the presence of Mg^2+^ with a *K*_*D*_ of ∼0.7 mM (**Figure 6A**). The pose of the backtracked nucleotide in our model is incompatible with backtracking by more than one nucleotide because the 3’OH of the backtracked nucleotide faces inwards. Finally, the observations that β’Q504R and β’K598W RNAPs displayed higher than expected intrinsic RNA cleavage activities (see below) further supports our inference that the backtracked nucleotide is at least fractionally localized in the E-site.

**Figure 6.**
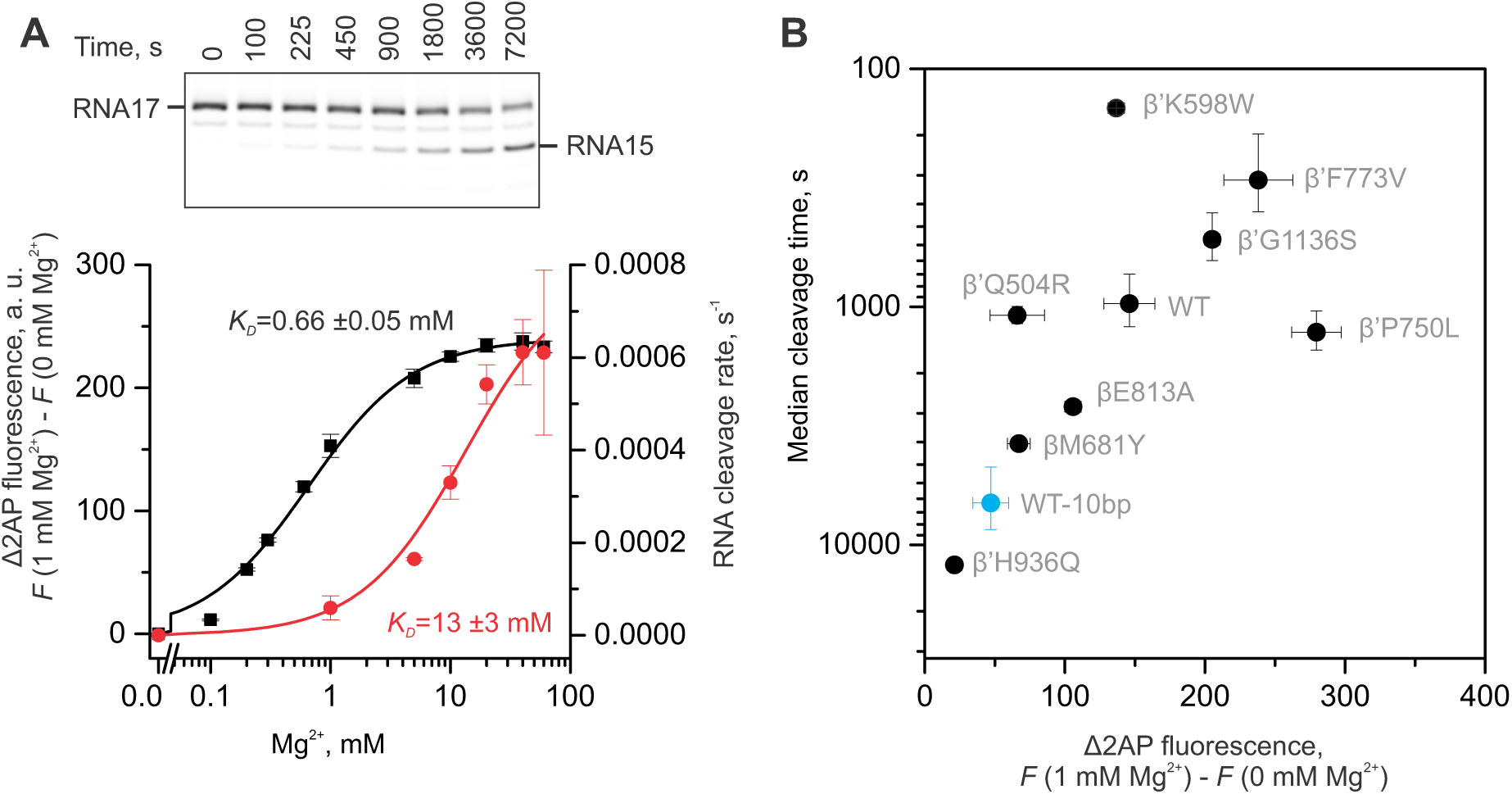
Intrinsic RNA cleavage activity of 2AP-TECs. **(A)** The dependence of the RNA cleavage activity (red) and fluorescence (black) of the 2AP-TEC on the Mg^2+^ concentration at pH 7.5. A representative gel of an RNA cleavage experiment performed at 10 mM Mg^2+^ is shown above the graph. **(B)** The effects of amino acid substitutions in the RNAP active site on the RNA cleavage activity and fluorescence of the 2AP-TECs. RNA cleavage experiments were performed at 1 mM Mg^2+^ pH 8.6 using 2AP-TECs with 11 bp (black circles) or 10 bp (blue circles) of RNA:DNA complementarity. The primary data are presented in **Supplementary Figure S4**. The fluorescence was measured at pH 7.5 (values are from **Figure 4A**). Error bars in (A) and (B) represent the range of duplicate measurements or standard deviations from multiple experiments.

### The intrinsic RNA cleavage activity positively correlates with the occupancy of the backtracked state

Most RNAPs possess a weak intrinsic RNA cleavage activity that is catalyzed by the same active site as the nucleotide addition reaction (Orlova *et al*, 1995). RNAPs most commonly excise two 3’-terminal nucleotides of the nascent RNA, though they can also cleave off one or three nucleotides under certain conditions. While it is uncertain if the intrinsic RNA cleavage activity has any physiological role in *E. coli*, this activity is commonly considered a hallmark of the backtracked state and has been shown to depend on the TL (Yuzenkova & Zenkin, 2010; Esyunina *et al*, 2015; Mishanina *et al*, 2017). The nascent RNA cleavage in 2AP-TEC was extremely slow (median cleavage time >10 000 s) under typical conditions used in our experiments (pH 7.5, 1 mM Mg^2+^). The slow RNA cleavage rate was partially due to a low Mg^2+^ concentration. Thus, the maximal RNA cleavage activity required a ∼20-fold higher Mg^2+^ concentration (apparent *K*_*D*_ ∼13 mM) than did the formation of the 1-nt backtracked state (apparent *K*_*D*_ ∼0.7 mM, **Figure 6A**). Apparently, the RNA cleavage reaction is accelerated by the binding of extra Mg^2+^ ions in addition to those needed for the stabilization of the backtracked state. However, even at 10 mM Mg^2+^ the rate of RNA cleavage by the wild-type RNAP (∼0.0003 s^-1^) was 6000-fold slower than the rate of backtracking (∼2 s^-1^), demonstrating that backtracking did not kinetically limit RNA cleavage in the 2AP-TEC. However, destabilization of the backtracked state by shortening the RNA:DNA complementarity to 10 bp inhibited the rate of RNA cleavage ∼10-fold (**Figure 6B**), suggesting that the cleavage rate is dependent on the occupancy of the backtracked state, i.e. backtracking thermodynamically limits the rate of RNA cleavage.

We further investigated the RNA cleavage activity of the 2AP-TECs assembled with variant RNAPs. To improve the accuracy and reproducibility of the measurements we increased the pH to 8.6, which increased the RNA cleavage rate ∼20-fold without significantly affecting the occupancy of the backtracked state. At the same time, we kept the Mg^2+^ concentration at 1 mM because (*i*) increasing the Mg^2+^ concentration increased the occupancy of the backtracked state and (*ii*) the variant 2AP-TECs differed in the apparent affinities for Mg^2+^ and the steepness of the concentration dependence curves (**Supplementary Figure S3**). Overall, the RNA cleavage rates positively correlated with the occupancy of the backtracked state, as expected. However, β´P750L displayed lower, whereas β´Q504R and β´K598W markedly higher, RNA cleavage activities than anticipated based on the backtracked state occupancy (**Figure 6B**).

The β’P750L can only affect the cleavage rate allosterically because it is located more than 20Å away from the backtracked nucleotide and the scissile bond (**Figure 5A**). Presumably, the conformation of the TH is slightly altered in the β’P750L TEC, leading to a modest reduction in the RNA cleavage rate. Indeed, β’P750L is located in the F-loop that has been shown to modulate the stability of the TH (Miropolskaya *et al*, 2009) and may therefore favor a TH conformation that is suboptimal for RNA cleavage. In contrast, the allosteric effects of β´Q504R and particularly β´K598W on the conformation of the TH are, in our view, relatively unlikely. Accordingly, the increase in RNA cleavage activities upon the β´Q504R and β´K598W substitutions that are located within the reach of the backtracked but not penultimate nucleotide (**Figure 5B**) suggests that the backtracked nucleotide is at least fractionally localized in the vicinity of β´Gln^504^ and β´Lys^598^.

### Backtracking does not limit GreA-facilitated RNA cleavage in the 2AP-TEC

The backtracked state has been traditionally associated with RNA proofreading by the excision of two or more 3’-terminal nucleotides from the backtracked RNA by Gre factors. In fact, sensitivity to Gre factors is routinely used to evaluate the propensity of the TEC to backtrack. To test whether GreA preferentially acted on the closed backtracked state, we pre-incubated the 2AP-TEC with and without Mg^2+^ prior to initiating the RNA cleavage reaction by addition of 8 µM of GreA. Unexpectedly, the rate of GreA cleavage was independent on the presence of Mg^2+^ during pre-incubation (**Figure 7**) despite the fact that the latter increased the occupancy of the backtracked state from ∼4 to ∼45% (**Figure 1C,D**) at the time of mixing with GreA. Irrespective of the exact mechanism behind the complex kinetics of GreA facilitated RNA cleavage (see below), the above experiments suggest that backtracking did not limit GreA-facilitated RNA cleavage in the 2AP-TEC. Otherwise, the preformed backtracked state in the 2AP-TEC supplemented with Mg^2+^ would cleave faster than the 2AP-TEC challenged with GreA without pre-incubation with Mg^2+^. This inference contrasts with the results of Turtola & Belogurov 2016 who reported that GreA-facilitated cleavage of the nascent RNA in the CMP extended 2AP-TEC was limited by backtracking (Turtola & Belogurov, 2016). The most likely reason for this difference is that the 2AP-TEC backtracks about 30 times faster (half-life ∼0.35 s, **Figure 2B**) than CMP-extended 2AP-TECs (median time ∼10 s; (Turtola & Belogurov, 2016)).

**Figure 7.**
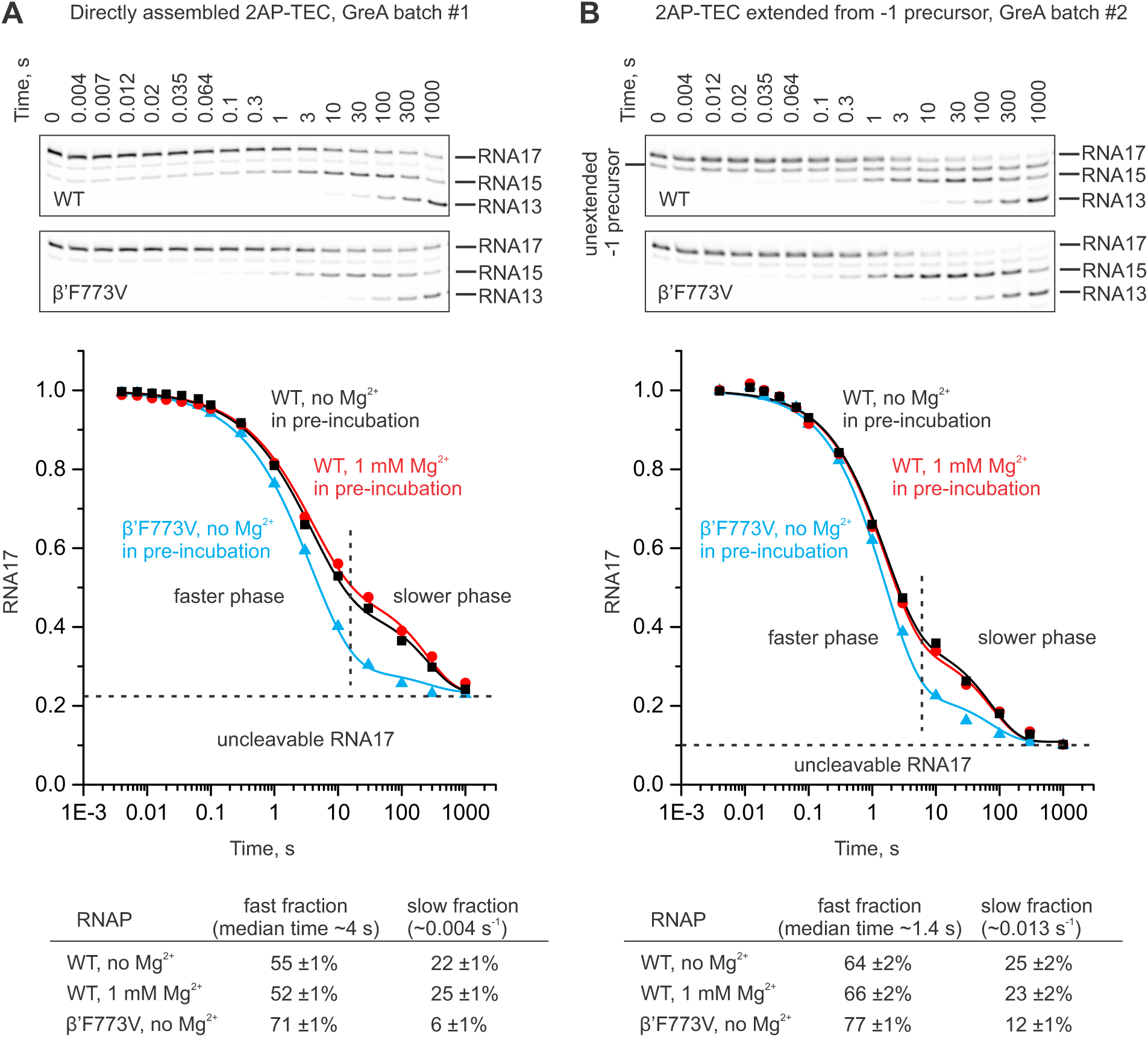
GreA facilitated cleavage of the nascent RNA in the 2AP-TEC. 2AP-TECs were directly assembled using a 17 nt RNA with a 2AP at the 3’ end **(A)** or prepared by the extension of a -1 precursor TEC with a 2AP triphosphate followed by gel filtration **(B)**. The graphs show the time courses of the RNA cleavage in 2AP-TECs in the presence of 8 µM GreA. Different batches of GreA were used in (A) and (B). The 2AP-TECs were assembled using the wild-type (black and red) or β’F773V (blue) RNAPs and pre-incubated with (black) or without (red and blue) 1 mM Mg^2+^ before mixing with an equal volume of 16 µM GreA. The final concentration of Mg^2+^ was 1 mM in all reactions. Lines are bestfits to the sum of the stretched exponential (corresponds to the faster phase) and exponential (corresponds to the slower phase) functions. Representative denaturing PAGE gels used to quantify the time courses of the RNA cleavage reactions are presented above the graphs. The bestfit values of the amplitudes of the fast and slow fraction of the 2AP-TECs are presented below the graphs. The fraction of the un-cleavable RNA and the rates of the fast and the slow phases were shared by the three data sets in each graph during the global fitting of the data (see the Methods section for details).

The time-course of the GreA-facilitated RNA cleavage (disappearance of RNA17) was well described by a sum of a stretched exponential function (corresponds to the faster phase), a single exponential function (corresponds to the slower phase), and a fraction of RNA17 that was resistant to cleavage (**Figure 7**). The kinetics of GreA-facilitated RNA cleavage displayed large variation (rates of both phases are approximately threefold slower in **Figure 7A** than in **Figure 7B**) depending on the batch of the GreA protein, even though we used a near saturating concentration of GreA. In addition, the fraction of the uncleaved RNA17 was larger when the 2AP-TEC was directly assembled than when the 2AP-TEC was obtained by extension of the -1 precursor TEC with the 2AP triphosphate. However, under identical conditions, GreA-facilitated cleavage in a β’F773V 2AP-TEC always displayed a larger amplitude of the fast phase than the wild-type 2AP-TEC, whereas the slow fraction was largely absent. The simplest interpretation is that the fast and slow phases correspond to the GreA-facilitated RNA cleavage activity of the 2AP-TECs backtracked by one and three nucleotides, respectively (**Figure 8A**). Whereas sequential 2+2 nt cleavage of the RNA apparently took place in both the wild-type and β’F773V 2AP-TECs, the slow phase of the disappearance of RNA17 (**Figure 7**, graphs) approximately corresponded to the appearance of the RNA13 (**Figure 7**, gel panels) that may partially originate from direct cleavage by four nucleotides. In other words, we suggest that the wild type 2AP-TEC contains a fraction that backtracked by more than one nucleotide (intrinsically present or induced by the binding of GreA – the cleavage factors have been shown to mobilize the backtracked RNA (Wang *et al*, 2009; Cheung & Cramer, 2011)), which results in the partial slowdown of the RNA cleavage reaction. At the same time, the β’F773V 2AP-TEC is resistant to multi-nt backtracking because it forms a very stable 1-nt backtracked state with the closed active site that inhibits further backtracking.

**Figure 8.**
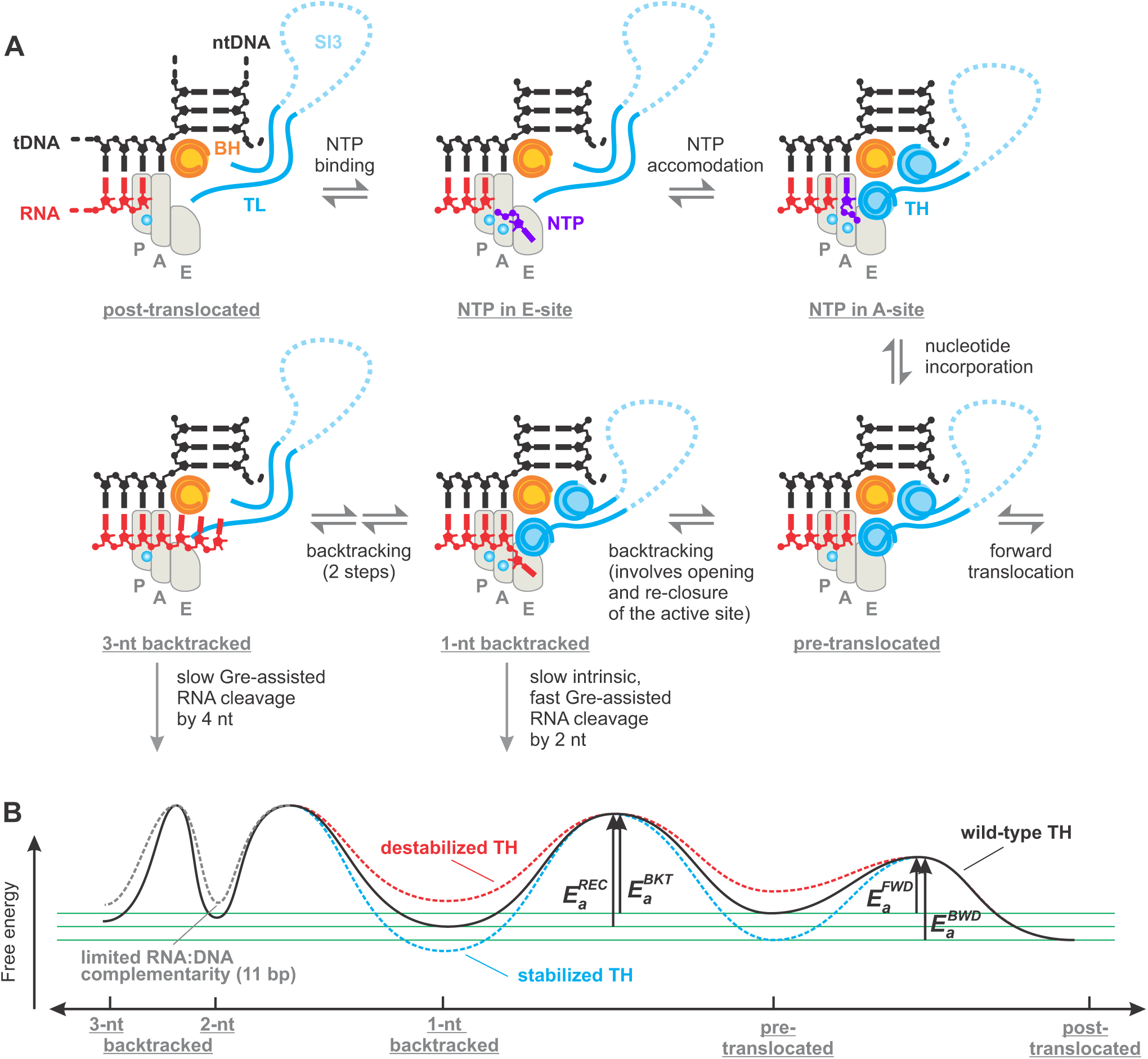
Backtracking and NTP substrate loading into the active site may follow the same route. **(A)** Schematics of the RNAP active site; only BH, TL/TH and the nucleic acids near the active site are shown. The areas corresponding to P-, A-and E-sites are outlined by light grey shapes. *Top row:* the triphosphate moiety of the substrate NTP governs binding to the E-site followed by rotation into the A-site where the complementarity to the template DNA is probed (71). The cognate NTP stabilizes the closed active site that positions the triphosphate moiety for efficient catalysis (1). *Bottom row:* following nucleotide incorporation RNAP completes the cycle by translocating forward but may occasionally backtrack. Backtracking by 1 nt places the backtracked nucleotide into the E-site; the backtracked and the penultimate nucleotides stabilize the closed active site. Further backtracking inhibits the closure of the active site. **(B)** The effect of the TH on the energy profile of the translocational equilibria. Green horizontal lines aid comparison of the energy minima. The heights of the activation energy barriers (*E*_*a*_) are not to scale (See **Supplementary Figure 7B**). *E*_*a*_ superscripts *REC, BKT, FWD* and *BWD* stand for the recovery, backtracking, forward and backward translocation, respectively. The energy profile corresponds to the 6AP-TEC: the relative depths of energy minima vary with the transcribed sequence.

## DISCUSSION

In this study we investigated the kinetic and structural properties of the simplest off-pathway state where RNAP is backtracked by 1-nt. Although backtracking was discovered over two decades ago and has immense physiological significance, several important questions have remained unanswered. First, most studies investigated the multi-nucleotide backtracking and had insufficient spatio-temporal resolution to directly visualize the entry into or recovery from a 1-nt backtracked state (Komissarova & Kashlev, 1997a; Depken *et al*, 2009; Ó Maoiléidigh *et al*, 2011; Dangkulwanich *et al*, 2013; Shaevitz *et al*, 2003). Accordingly, it remained uncertain how the kinetics of backtracking by 1-nt compares with that of the subsequent backtracking steps. Second, it is not known how the rate of recovery from the 1-nt backtracked state compares with the forward rates characteristic of processive elongation. Does backtracking by one nucleotide induce pausing or is the backtracked state in rapid equilibrium with the on-pathway states? Indeed, the Brownian ratchet model of RNAP translocation postulates that the same amount of energy is available for the forward translocation from the pre-to the post-translocated state and from the backtracked to the pre-translocated state. Accordingly, there is no a priori indication that the energy barrier between the 1-nt backtracked and the pre-translocated states is uniformly high enough to equate short backtracking with pausing. Third, it is uncertain if the 1-nt backtracked state is stabilized exclusively by contacts between the backtracked nucleotide and the open active site of RNAP, as suggested by crystallographic evidence (Wang *et al*, 2009; Sekine *et al*, 2015), or if a certain conformer of the TL plays a role in maintaining the stability of the backtracked state (Mishanina *et al*, 2017).

### Active site closure stabilizes the 1-nt backtracked state

The extent of backtracking can be precisely limited to one nucleotide by assembling the TEC on nucleic acid scaffolds and limiting the RNA:DNA complementarity to 11 base pairs. A far bigger challenge was to design a TEC that fractionally backtracks. The predominantly backtracked TECs are commonly designed by using an RNA primer with a 3’ end mismatch against the template DNA. However, the latter setup precludes the accurate estimation of the recovery kinetics because the 3’ end mismatch significantly slows down the incorporation of nucleotides by a recovered post-translocated TEC. We overcame these limitations by designing a TEC with an unusual 2AP-dT pair at the downstream edge of the RNA-DNA hybrid (**Figure 1A,B**). An alternative setup with inosine in the template DNA was also considered. However, the 2AP setup was preferred because 2AP (*i*) forms a Watson-Crick base pair with thymine, in contrast to a wobble base pair formed by inosine, (*ii*) preserves more than half of the catalytic activity of the post-translocated TEC with an adenine at the 3’ end **(Figure 1D and Supplementary Table S1**), thereby allowing accurate measurements of the recovery from the backtracked state, and (*iii*) additionally allows assessing the conformation of the RNA 3’ end by monitoring the 2AP fluorescence.

We show that the TEC bearing the 2AP in the nascent RNA 3´ end undergoes a fractional transition into an inactivated state upon the addition of Mg^2+^. We then demonstrate that the inactivated state bears the hallmarks of the backtracked state. First, formation of the backtracked state leads to an increase in the fluorescence of the 2AP nucleotide at the RNA 3’ end consistent with its unstacking from the penultimate guanine base as a result of backtracking. Second, the fraction of the inactivated state and the fluorescence of 2AP-TEC were reduced by the RNA:DNA and the DNA:DNA mismatches that disfavor the backward movement of the transcription bubble from the post-translocated to the 1-nt backtracked state.

We next examined the occupancies of the backtracked state in a set of active site variants. We found that the amino acid substitutions that stabilize the closed active site markedly increased the occupancies of the backtracked state, whereas substitutions of the residues that as a part of the TH interact with the penultimate RNA nucleotide in the substrate site (aka the A site) decreased the occupancies of the backtracked state. However, we noted that substitutions can potentially affect the occupancy of the backtracked state by solely biasing the equilibrium between the pre-and post-translocated states. Further analyses revealed that amino acid substitutions that stabilize the closed active site markedly reduce the recovery rate from the backtracked state whereas substitutions that destabilize the closed active site increase the recovery rate. The latter results strongly suggest that the closed active site directly stabilizes the backtracked state.

### Structural model of the 1-nt backtracked state with the closed active site

Next, we carried out modeling experiments aimed at testing whether the closed active site can accommodate the backtracked nucleotide because the crystal structures of the 1-nt backtracked state available at the time of writing were incompatible with the closed active site. We found that the backtracked purine nucleotide does not readily fit into the interior of the closed active site but can be accommodated in the secondary channel area known as the E-site. Next, the effects of the amino acid substitutions on the occupancy and stability of the backtracked state did not pinpoint the location of the backtracked nucleotide because interplay between direct and allosteric effects could be anticipated in most cases. Thus, each substitution can affect the stability of the backtracked state by altering the stability of the TH or by directly modifying the contacts with the backtracked or penultimate RNA nucleotide. However, the analysis of the correlation between the intrinsic RNA cleavage activity and the occupancy of the backtracked state in variant RNAPs supported the E-site localization of the backtracked nucleotide. Specifically, two out of three outliers in the correlation graph were the amino acid substitutions in the E-site (β´Q504R and β´K598W, **Figure 5B**). In addition, Zenkin et al observed that AMPCPP, the non-hydrolysable analogue of ATP, inhibited the intrinsic RNA cleavage activity in the TEC with a mismatch between the RNA 3’ end and the template DNA (Zenkin *et al*, 2006). Given that the AMPCPP was not complementary to the post-translocated state of the mismatched TEC and could only bind to the E-site, they suggested that the inhibition was due to the competition between the AMPCPP and the mismatched RNA 3’ end for binding to the E-site. These results provide additional support for the E-site localization of the backtracked nucleotide. However, in our model the backtracked nucleobase is not positioned to directly coordinate Mg^2+^ as suggested by Zenkin et al (Zenkin *et al*, 2006) and facilitates the RNA cleavage allosterically by stabilizing the backtracked state and modulating the conformation of the penultimate RNA nucleotide. Our model also differs from the model proposed by Sosunova et al where the nucleobase of the backtracked nucleotide in the E-site faces into the interior of the active site (Sosunova *et al*, 2013) (**Supplementary Figure S5**). While the pose of the backtracked nucleotide proposed by Sosunova et al is compatible with the closed active site, it does not explain the markedly elevated RNA cleavage activity of β´K598W RNAP. That said, it cannot be ruled out that poses of the backtracked nucleotide with the nucleobase facing outward (our model) and inward (Sosunova *et al*, 2013) coexist and interconvert.

Our data show that formation of the 1-nt backtracked state is dependent on the presence of millimolar concentration of Mg^2+^ but do not pinpoint the exact positions of the Mg^2+^ ions in the structure. Instead, we suggest approximate locations of the Mg^2+^ ions based on their typical positions in the active site of the multisubunit RNAPs. All Mg^2+^ ions coordinated by the RNAP active site bind to a cluster of five acidic residues consisting of the invariant β’ Asp triad (β’Asp^460^, Asp^462^, Asp^464^ in *Eco* RNAP) and an acidic pair contributed by the β subunit (βGlu^813^ and Asp^814^ in *Eco* RNAP). With the rare exceptions, RNAP structures contain a high-affinity Mg^2+^ ion (MG1) bound to the Asp triad. In the post-translocated state, MG1 binds and activates the 3’OH group of the nascent RNA for the nucleophilic attack on the α phosphate of the NTP substrate. In the pre-translocated state, MG1 bridges the Asp triad with the phosphate of the 3’ terminal RNA nucleotide. We suggest that MG1 similarly bridges the Asp triad with the penultimate RNA phosphate in the 1-nt backtracked state (**Figure 5B, Supplementary Figure S6**). RNAP structures with the phosphate-bearing ligands in the active site often contain an additional Mg^2+^ ion (MG2) coordinated by the β acidic pair (either directly or through water molecules) and one or two residues from the β’ Asp triad. In some cases, MG2 plays only a structural role by bridging the phosphate groups of the ligands with the cluster of acidic residues in the RNAP active site (Artsimovitch *et al*, 2004; Vassylyev *et al*, 2005). In the other cases, MG2 also plays a catalytic role: it stabilizes the pentacovalent transition state and assists the leaving of pyrophosphate during nucleotide incorporation (Wang *et al*, 2006; Vassylyev *et al*, 2007b) or activates a water molecule for a nucleophilic attack on the RNA phosphate during the cleavage of the nascent RNA in the RNAP active site (Sosunov *et al*, 2003; Sosunova *et al*, 2013). We suggest that MG2 binds close to βGlu^813^ where it bridges the hydroxyl groups of the backtracked nucleotide with the cluster of acidic residues in the RNAP active site, but cannot position and activate the water molecule for an attack on the RNA phosphate (**Figure 5B, Supplementary Figure S6**). In contrast, β’Q504R and β’K598W substitutions modestly remodel the active site (via contact with the nucleobase in the latter case) and facilitate the fractional binding of MG2 in an alternative location (presumably closer to βAsp^814^) where it can participate in water activation thereby increasing the RNA cleavage activity of the backtracked state (**Supplementary Figure S6**). Importantly, the requirement for MG2 for positioning of the backtracked nucleotide possibly explains the structural differences between the backtracked state in our system (**Figure 5B**) and the backtracked states crystalized in the absence of the added Mg^2+^ (Wang *et al*, 2009; Sekine *et al*, 2015) (**Figure 5C**).

### Backtracking by 1 nt resembles the reversal of the NTP substrate loading

We note that the reversible migration of the RNA 3’ NMP from the A-site to the E-site during entry and recovery from the 1-nt backtracked state are reminiscent of the proposed pathway for NTP loading and exit from the active site (Batada *et al*, 2004; Westover *et al*, 2004), except that the 3’NMP is covalently attached to the nascent RNA (**Figure 8A**). While the NTP in the E-site did not stabilize the TH to a measurable extent (Westover *et al*, 2004; Wang *et al*, 2006), the backtracked nucleotide can stabilize the TH indirectly via contacts with the penultimate nucleotide in the A-site (e.g. β’M932 -nucleobase interactions). Indeed, perturbing these contacts by the β’M932L or β´N458D substitutions reduced the occupancy of the backtracked state (**Figure 4A**). In addition, binding of the 3’NMP to the E-site positions its phosphate group for interactions with β’Gln^929^ and β’His^936^ in the TH and these interactions may contribute to the stabilization of the closed active site. Consistently, the β’Q929A, β’H936A and β’H936Q substitutions led to a decrease in the occupancy of the backtracked state (**Figure 4A,B**). Moreover, we demonstrated that the 1-nt backtracked state was at least 2.5-fold less stable in β’H936Q than in the wild-type RNAP (**Figure 4C**).

Interestingly, the variant RNAPs that display a higher fraction of the backtracked state in our study have been previously characterized as pause-resistant (Svetlov *et al*, 2007; Malinen *et al*, 2014) and were presumed to backtrack less than the wild-type RNAP (Bar-Nahum *et al*, 2005; Nedialkov *et al*, 2013). There are three reasons for this superficial inconsistency. First, the stabilized 1-nt backtracked state inhibits further backtracking as suggested by the GreA-facilitated RNA cleavage experiments shown in **Figure 7,** thereby preventing long-term arrests during processive transcript elongation. Second, the stabilized closed active site increases the depth of the energy minima corresponding to the pre-translocated and the backtracked states, thereby also increasing the activation energy barrier separating the two states (**Figure 8B**). Third, the kinetic competition between the backtracking and nucleotide addition rather than the thermodynamic stability of the backtracked state determines the probability of entering the backtracked state during processive transcript elongation (Dangkulwanich *et al*, 2013). The variant RNAPs with the stabilized closed active site have higher catalytic efficiencies than the wild-type RNAP and are therefore less likely to backtrack at sub-saturating concentration of NTPs often employed in *in vitro* transcription experiments (Svetlov *et al*, 2007).

We suggest that in our systems (2AP-and 6AP-TECs) backtracking originates from the pre-translocated state where it directly competes with the forward translocation: opening of the active site reduces the affinity of the A-site for the RNA 3’ end (Feig & Burton, 2010; Malinen *et al*, 2012) allowing the latter to move forward into the P-site or backward into the E-site (**Figure 8**). Our data suggest that RNAP has about 1-4% chance to backtrack following the incorporation of AMP at the sequence position corresponding to the 6AP-TEC during the processive transcript elongation (**Supplementary Figure S7**). However, the backtracking rate and the propensity to backtrack following the addition of CMP at the next sequence position is ∼30 fold lower (see Gre section in the Results section). In addition, backtracking may originate from other translocational states such as an elemental pause at positions where these states get populated during the transcript elongation (Artsimovitch & Landick, 2000; Kang *et al*, 2018). We thus caution against relating the inferred backtracking propensity of 6AP-TEC to the average backtracking propensity. Instead, our data provide an informative example of the backtracking propensity attributable to the direct competition between the backtracking and the forward translocation following the incorporation of a cognate substrate at a non-pause position.

In summary, our data depict the predominant fraction of a 1-nt backtracked TEC as a distinct state that resembles the pre-translocated state with a closed active site and the backtracked nucleotide in the E-site but is different from states where RNAP backtracks by many nucleotides. Recovery from this state likely relies on the same conformational transitions that mediate substrate loading and thermally-driven translocation during the on-pathway elongation accounting for a relatively fast escape rate (median lifetime ∼2 s). Our results possibly provide the mechanistic basis for the inferences of the single-molecule study by Dangkulwanich et al who concluded that the first backtracking step is distinct from subsequent steps (Dangkulwanich *et al*, 2013), despite we reached opposite conclusions about the stability of the 1-nt backtracked state in the altered RNAPs with the stabilized closed active site. Interestingly, the lifetimes of the backtracking events measured here generally parallel the short sequence-specific pauses observed in single-molecule experiments (Herbert *et al*, 2006; Shundrovsky *et al*, 2004), some of which may be caused by short backtracking (Galburt *et al*, 2007; Depken *et al*, 2009; Ó Maoiléidigh *et al*, 2011). These characteristics suggest that the closed 1-nt backtracked states both play an important role in the control of the sequence specific elongation rate and serve as failsafe checkpoints preventing RNAP from sliding further backwards into the long-backtracked states that lead to prolonged inactivation or permanent arrest.

## MATERIALS AND METHODS

### Reagents and oligonucleotides

DNA and RNA oligonucleotides were purchased from Eurofins Genomics GmbH (Ebersberg, Germany) and IBA Biotech (Göttingen, Germany). DNA oligonucleotides and RNA primers are listed in **Supplementary table S1**. NTPs were from Jena Bioscience (Jena, Germany), 2AP-5´TP was from TriLink Biotechnologies (San Diego, USA). Streptolydigin was purchased from Sourcon-Padena (Tübingen, Germany).

### Proteins

RNAPs were expressed in *E. coli* Xjb(DE3) (Zymo Research, Irvine, CA, USA) or T7 Express lysY/I^q^ (New England Biolabs, Ipswich, MA, USA) and purified by Ni-, heparin and Q-sepharose chromatography as described previously (Svetlov & Artsimovitch, 2015), dialyzed against the storage buffer (50% glycerol, 20 mM Tris-HCl pH 7.9, 150 mM NaCl, 0.1 mM EDTA, 0.1 mM DTT) and stored at -20°C. The GreA protein containing a C-terminal His^6^-tag was purified by immobilized-metal affinity chromatography followed by gel filtration. Plasmids used for protein expression are listed in **Supplementary Table S2**.

### TEC assembly

TECs (1 µM) were assembled by a procedure developed by Komissarova et al. (Komissarova *et al*, 2003). An RNA primer (final 1 µM) was annealed to the template DNA (final 1.4 µM), incubated with RNAP (1.5 µM) for 10 min, and then with the non-template DNA (2 µM) for 20 min at 25 °C. The assembly was carried out in TB0 buffer (40 mM HEPES-KOH pH 7.5, 80 mM KCl, 5% glycerol, 0.1 mM EDTA, and 0.1 mM DTT) or in TB1 buffer (TB0 supplemented with 1 mM MgCl^2^). TEC17 with the 2AP in the RNA 3´ end (2AP-TEC) was either directly assembled using a 17 nt long RNA with 2AP or formed by extension of the assembled TEC16 by RNAP. In the latter case, the assembled TEC16 was incubated with 50 µM 2AP-triphosphate and 5 µM GTP for 2 minutes. GTP was included to re-extend the minor fraction of RNAs that got cleaved by 2-nt during extension. The 2AP-TEC was then either diluted fivefold for use in CMP incorporation experiments or purified from Mg^2+^ and NTPs for use in the other assays. Removal of Mg^2+^ and NTPs was carried out using Zeba Spin Desalting Columns 40K MWCO (Pierce Biotechnology, Rockford, USA) pre-equilibrated with TB0. The nucleic acid scaffolds used for assembling the TECs are presented in **Supplementary Figure S1**.

### Time-resolved nucleotide addition and GreA-facilitated RNA cleavage measurements

The measurements were performed in an RQF 3 quench-flow instrument (KinTek Corporation, Austin, TX, USA). The reaction was initiated by the rapid mixing of 14 µl of 0.2 µM 2AP-TEC with 14 µl of 400 µM CTP or 16 µM GreA. The 2AP-TEC solution was prepared either in TB0 or TB1 buffer. In the former case, CTP and GreA solutions were prepared in TB2 buffer (TB0 supplemented with 2mM MgCl^2^), whereas in the latter case CTP and GreA solutions were prepared in TB1 buffer. GreA solutions additionally contained 300 mM KCl. The reaction was allowed to proceed for 0.004–1000 s at 25 °C, followed by quenching with 86 µl of 0.5 M HCl and neutralization by 171 µl of neutralizing buffer (94% formamide, 290 mM Tris base, 13 mM Li^4^-EDTA, 0.2% Orange G). RNAs were separated on 16% denaturing polyacrylamide gels and visualized with an Odyssey Infrared Imager (Li-Cor Biosciences, Lincoln, NE, USA); band intensities were quantified using the ImageJ software (Abramoff *et al*, 2004).

### Time resolved fluorescence measurements

Measurements were performed in an Applied Photophysics (Leatherhead, UK) SX.18MV stopped-flow instrument at 25 °C. The 2AP fluorescence was excited at 320 nm and the emitted light was collected through a 375 nm longpass filter. At least three individual traces were averaged for each reported curve. The experiments reporting the increase in fluorescence were initiated by mixing 60 µl of 0.1 µM 2AP-TEC in TB0 buffer with 60 µl of TB2 buffer. The experiments reporting the decrease in fluorescence were initiated by mixing 60 µl of 0.1 µM 2AP-TEC in TB1 buffer with 60 µl of 400 µM CTP solution in TB1 buffer.

### Equilibrium fluorescence measurements

Equilibrium fluorescence levels were determined by continuously recording the light emission at 380 nm (excitation at 320 nm) with an LS-55 spectrofluorometer (PerkinElmer, Waltham, MA, USA) in a 16.160-F/Q/10 quartz cuvette (Starna) at 25 °C. The 2AP-TECs were assembled and diluted to 0.2 µM in TB0 buffer. The Mg^2+^-free fluorescence levels were recorded followed by the addition of MgCl^2^ and other regents (CTP, Stl, where indicated). The fluorescence was allowed to equilibrate for up to two minutes between each successive addition. The fluorescence was corrected for the fraction of the active TEC in each preparation that was determined by the incubation of 2AP-TEC with 200 µM CTP for 2 min or with 8 µM GreA for 10 min. Following the addition of CTP or GreA, 4 µl aliquots were withdrawn from the cuvette, quenched with 6 µl of the gel loading buffer (94% formamide, 20 mM Li^4^-EDTA and 0.2% Orange G) and analyzed by PAGE as described above.

### Analyses of the intrinsic RNA cleavage activity of 2AP-TECs

In the experiments presented in **Figure 6A** (bottom, dark yellow circles), the rates of RNA cleavage were determined by monitoring the decrease in fluorescence after the addition of 1-60 mM MgCl^2^ to the 2AP-TEC in TB0 buffer. The 2AP fluorescence emission was recorded at 380 nm (excitation at 320 nm) with an LS-55 spectrofluorometer (PerkinElmer, Waltham, MA, USA) in a 16.160-F/Q/10 quartz cuvette (Starna) at 25 °C. In these experiments, the initial increase in fluorescence corresponding to the formation of the backtracked state occurs at the timescale of seconds and was not monitored in the manual-mixing experiment. In contrast, the subsequent RNA cleavage reaction led to a decrease in fluorescence and occurred at the timescale of minutes to hours. The fluorescence assay produced temporary dense datasets thereby allowing relatively rapid measurements of the very slow cleavage rates at a low Mg^2+^ concentration. We further confirmed that the RNA cleavage rate inferred from the fluorescence measurements at 10 mM Mg^2+^ matched the rate inferred from the gel based assay (**Figure 6A**, *top*). The latter assay was performed as described below except that TB0 buffer was used in place of TB0-8.6 buffer and the reaction was initiated by the addition of 10 mM MgCl^2^.

In the experiments presented in **Figure 6B,** TEC16 containing a non-template DNA with a biotin tag at the 5’ end was immobilized on magnetic streptavidin beads (Dynabeads® MyOne(tm) Streptavidin C1, Invitrogen), extended into 2AP-TEC by the addition of 50 µM of 2AP-triphosphate (5 µM of GTP was also included to re-extend the minor fraction of RNAs that got cleaved by 2 nt during extension) for 2 minutes in TB1 buffer, washed three times with TB0 buffer and suspended in TB0-8.6 buffer (40 mM TAPS pH 8.6, 80 mM KCl, 5% glycerol, 0.1 mM EDTA, and 0.1 mM DTT). Cleavage reactions were initiated by adding 1 mM of MgCl^2^, and stopped at indicated times by withdrawing 8 µl aliquots and mixing them with 12 µl of the gel loading buffer. The progress of the reaction was then evaluated by separating the product RNAs in the denaturing PAGE and quantifying the individual bands.

### Data analyses

The time courses of CMP incorporation were fitted to a sum of exponential (models fast phase) and stretched exponential (models slow phase) functions. The same approach was used to fit the time courses of the GreA facilitated cleavage of the nascent RNA, but the stretched exponential component corresponded to the fast phase, whereas the exponential component corresponded to the slow phase. In addition, the time courses of the GreA facilitated cleavage of the nascent RNA by the wild-type (pre-incubated with and without Mg^2+^) and β’F773V RNAP (pre-incubated without Mg^2+^) were analyzed globally with the shared rate parameters and stretching parameters for all three datasets, but with unique parameters describing the fractions of the fast and slow phase for each dataset. The stretched exponential function was also used to fit the decrease in the fluorescence upon CMP incorporation and time courses of the intrinsic cleavage of the nascent RNA. The median reaction times were calculated as in (Turtola & Belogurov, 2016). We extensively used the stretched exponential function in the analyses because it is the simplest analytical function that robustly describes the time courses of the intrinsic (this work) and GreA facilitated (this work and (Turtola & Belogurov, 2016)) cleavage of the nascent RNA as well as the recovery kinetics from the backtracked state (this work). At the same time, these relatively slow processes in transcription are, in most cases, poorly described by the single and double exponential functions (Turtola & Belogurov, 2016).

### Modeling the backtracked state with the closed active site

To generate a model of the backtracked state we selected the crystal structure of a pyrophosphate bound initially transcribing complex (ITC) of the *Eco* RNAP (PDB ID 5IPL) (Liu *et al*, 2016) as a starting model (pyrophosphate was omitted from the structure). While our biochemical experiments were done with the TEC rather than ITC, the downstream parts of the transcription bubble, including the active site, are largely identical in the TEC and ITC as judged by a comparison of the *Thermus thermophilus* TEC (Vassylyev *et al*, 2007a, 2007b) and ITC structures (Zhang *et al*, 2012; Bae *et al*, 2015; Basu *et al*, 2014). Also, the ITC structures were the only structures of the *Eco* RNAP in the pre-translocated state with the closed active site available at the time of writing. Out of several *Eco* ITC structures (Zuo & Steitz, 2015; Liu *et al*, 2016) the one containing pyrophosphate appeared the most suitable, as it was inferred from the highest resolution electron density map and, as our trials showed, featured a helical TL conformation that allowed for the least constrained accommodation of the phosphate of the backtracked nucleotide. Given that in the starting model, the 3’NMP is disjoined from the rest of the RNA (due to an equilibrium between nucleotide addition and pyrophosphorolysis or intrinsic RNA cleavage in the crystals) we first remodeled the 3’NMP to generate a continuous pre-translocated RNA. The position of the 3’NMP was guided by the structures of the pre-translocated ITC that feature the continuous RNA densities (PDB ID 5IPM, 4Q5S), Watson-Crick base pairing with the template DNA, and the permitted nucleic acid geometries. The pre-translocated model was then extended by attaching a 2AP nucleoside monophosphate to the pre-translocated RNA 3’ end and the torsion angles around the phosphate and the C5’ of the backtracked nucleotide as well as the C3’ of the penultimate nucleotide were manipulated to search for the spatially feasible poses of the backtracked nucleotide. We found that the position of the phosphate of the backtracked nucleotide is almost uniquely defined due to steric hindrance imposed by the TH, the β’N458, the ribose and the phosphate of the penultimate nucleotide. In contrast, the sugar and the nucleobase of the backtracked nucleotide could be accommodated in several distinct conformations within an elongated opening that stretches along the N-terminal helix of the TL and connects the interior of the active site with the external milieu (see the Discussion for details). We also note that additional interactions between the TH with the 2’OH group of the penultimate nucleotide can be achieved by adjusting the conformation of the β’Q929 sidechain. The atomic coordinates of the model of the 1-nt backtracked state complex are provided as **Supplementary Dataset 1** (BKT-ITC.pdb)

## AKNOWLEDGEMENTS

The authors would like to thank Dr. Irina Artsimovitch and Dr. Robert Landick for critically reading the manuscript; Dr. Irina Artsimovitch for providing expression plasmids and the instruments for the anisotropy measurements at the Ohio State University; Dr. Thadée Grocholski and Dr. Anssi Malinen for assistance with protein purification and cloning. Essential equipment was contributed by the Walter and Lisi Wahl Foundation.

---

**Supplementary Figure S1.**
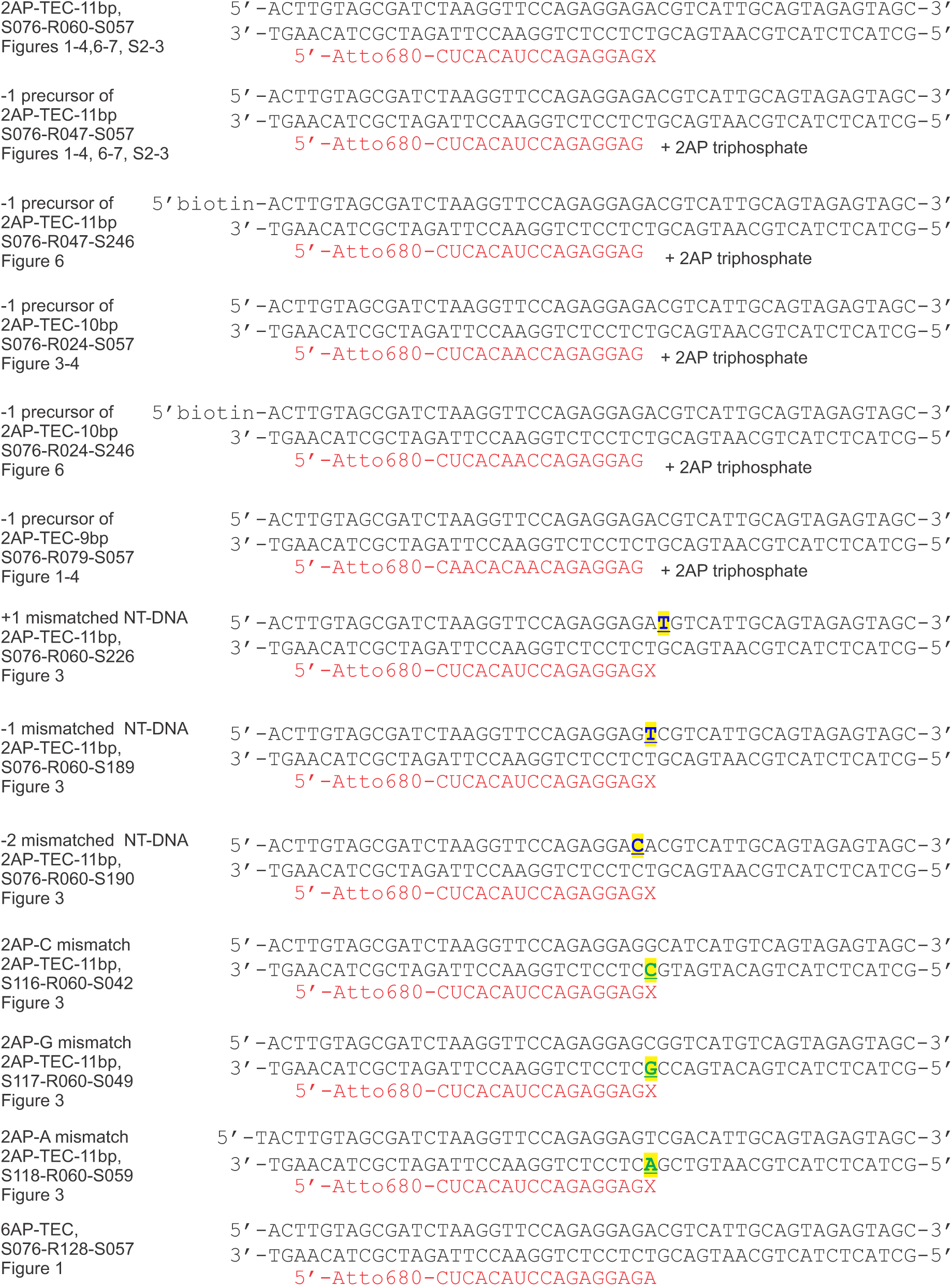
TEC systems employed in this work. X = 2AP, DNA:DNA (blue) and DNA:RNA (green) mismatches are underlined and highlighted in yellow. The identifiers for oligonucleotides (internal naming used in the laboratory) used for scaffold assembly are indicated alongside the names of the TECs.

**Supplementary Figure S2.**
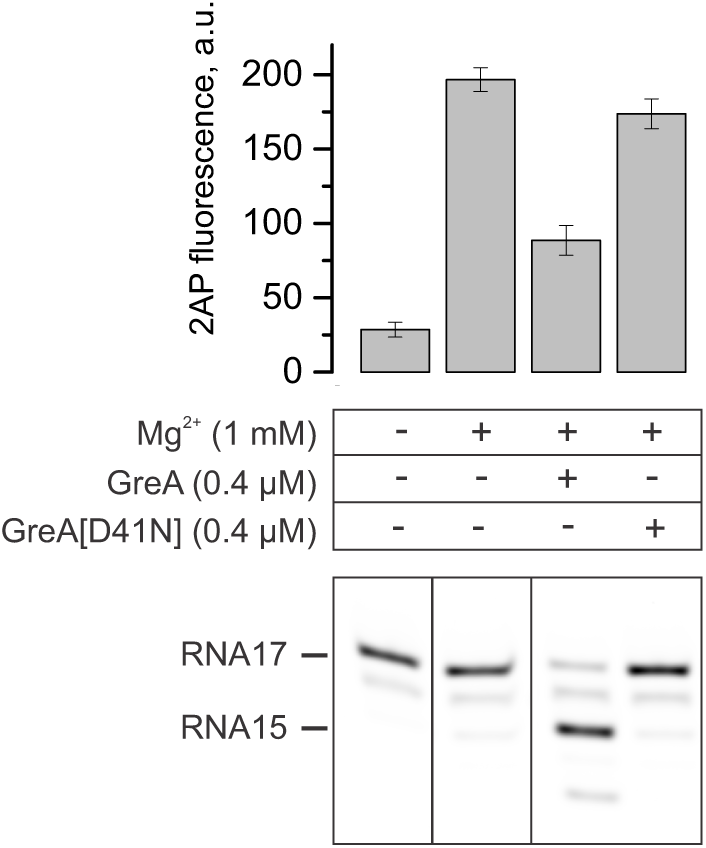
Endonucleolytic cleavage of the nascent RNA in the 2AP-TEC decreases the 2AP fluorescence. A schematic of the 2AP-TEC used in these experiments is shown in **Figure 1B**. The addition of Mg^2+^ to a Mg^2+^-free 2AP-TEC increases the fluorescence, further addition of the wild-type GreA (but not the cleavage-deficient GreA[D41N]) decreases the fluorescence more than twofold. Error bars show the range of duplicate measurements. Adenaturing PAGE gel of the RNA in the 2AP-TECs is shown below the bar graph. The gel panels were spliced from the same gel.

**Supplementary Figure S3.**
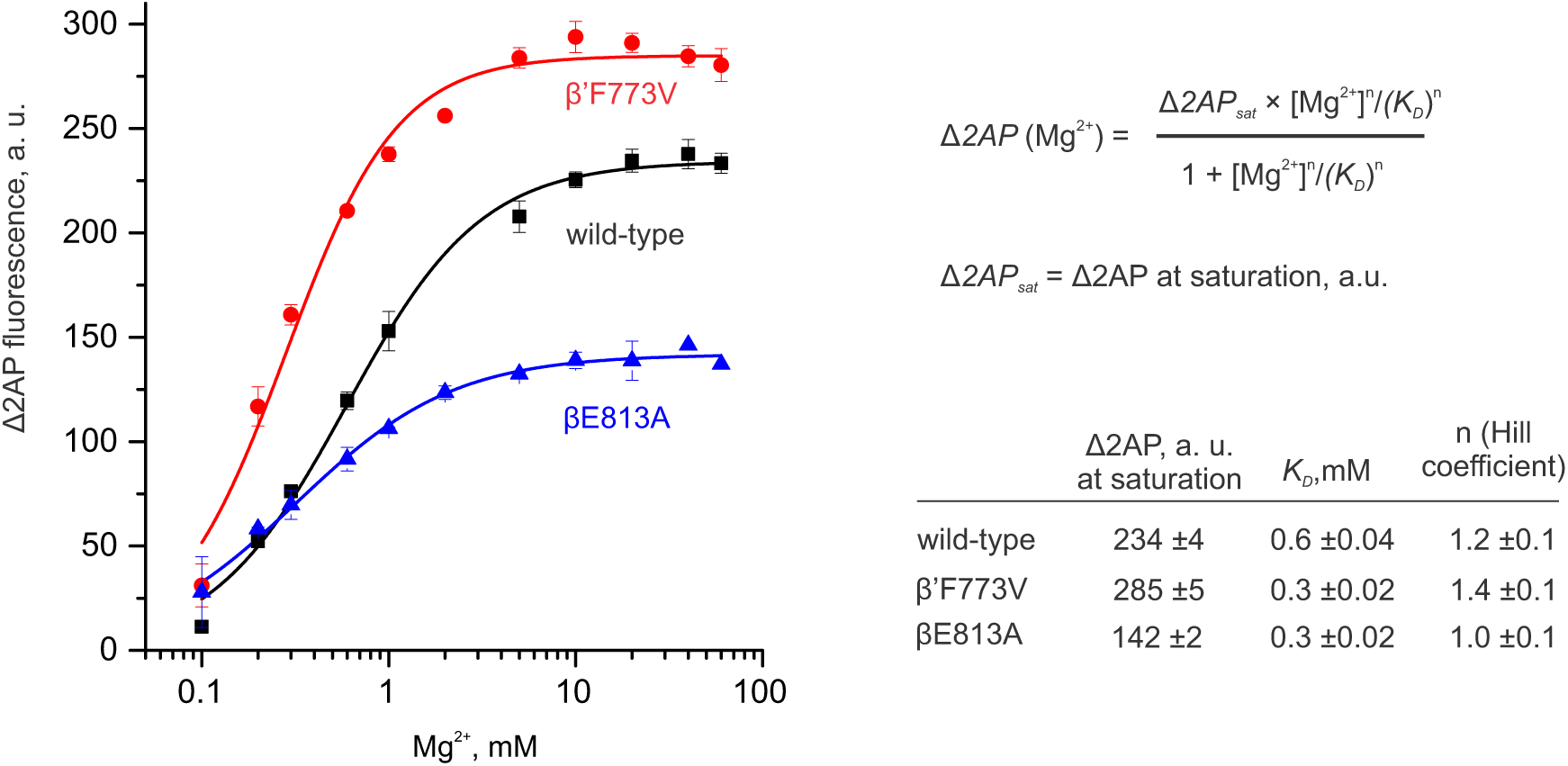
The dependence of the 2AP-TEC fluorescence on the Mg^2+^ concentration at pH 7.5. Error bars in the graph represent the range of duplicate measurements or standard deviations from multiple experiments. The errors in the table are SDs estimated by the nonlinear regression analysis of the data in the graph using the Hill equation (*Top right*). The Adair equation with two sites is mechanistically more appropriate for the wild-type and β’F773V data, but fitting the data using the Adair equation resulted in poorly constrained parameters. The Hill coefficients of the βE813A and the wild-type RNAP are not decisively different considering SDs, but the data are overall consistent with the hypothesis that the backtracked state is stabilized by a single Mg^2+^ ion bound to the Asp triad in the βE813A RNAP, whereas more Mg^2+^ ions participate in the stabilization of the backtracked state in the wild-type and the β’F773V RNAPs (see the main text **Figure 5B** for details).

**Supplementary Figure S4.**
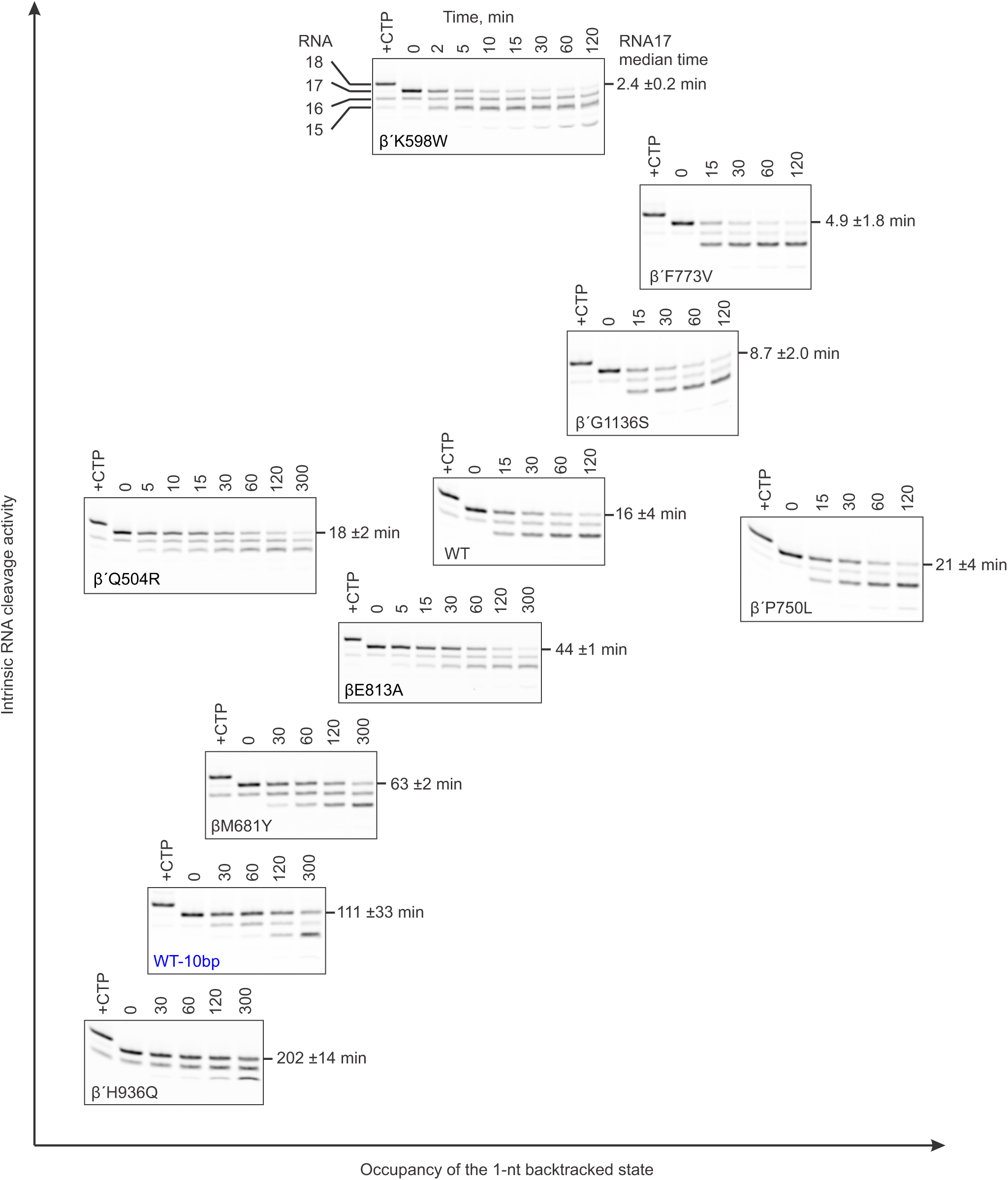
Primary data for the main text figure 6B. The denaturing PAGE gels show the results of the RNA cleavage experiments and are arranged to approximately follow the correlation graph in the main text **Figure 6B.** Note that the median cleavage times (values on the right from each gel panel) were calculated for the disappearance of RNA17 and do not differentiate between cleavage by 2 nt catalyzed by the backtracked state (prevalent mode in the wild-type and most altered RNAPs) and cleavage by 1 nt catalyzed by the pre-translocated state (prevalent mode in β’H936Q).

**Supplementary Figure S5.**
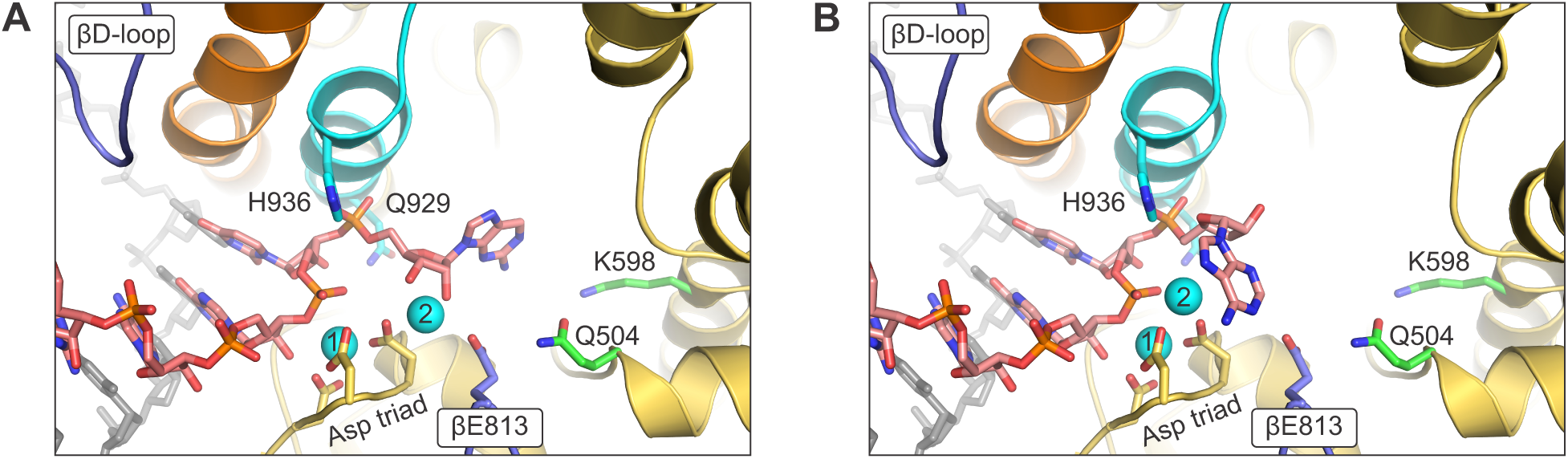
A side-by-side comparison of the backtracked nucleotide in our model. **(A)** with the pose proposed by Sosunova et al (81) **(B)**. Only selected regions of β subunit are shown. Mg^2+^ ions are depicted as cyan spheres and numbered according to their relative affinity: high-affinity MG1 and lower affinity MG2. Other colors are as in the main text Figure 5A. The atomic coordinates published by Sosunova et al comprise selected loops of the active site of the yeast Pol II with the backtracked nucleotide positioned based on the biochemical analysis of *Eco* RNAP. To facilitate comparison, we constructed *Eco* RNAP model with the backtracked nucleotide in a conformation proposed by Sosunova et al instead of using coordinates of their “hybrid” model. To prepare the image in (B) we used our model as a starting structure, changed the nucleobase of the backtracked nucleotide to adenine, altered the overall pose of the backtracked nucleotide and relocated MG2 to match their positions in the model published by Sosunova et al (81). Note that we positioned MG2 far away from the penultimate phosphate to render the backtracked state inactive in RNA cleavage, whereas Sosunova et al positioned MG2 close to the penultimate phosphate to model a cleavage-proficient state (See **Supplementary Figure S6** for details). Figure was prepared using PyMOL Molecular Graphics System, Version 1.8.6.0 Schrodinger, LLC.

**Supplementary Figure S6.**
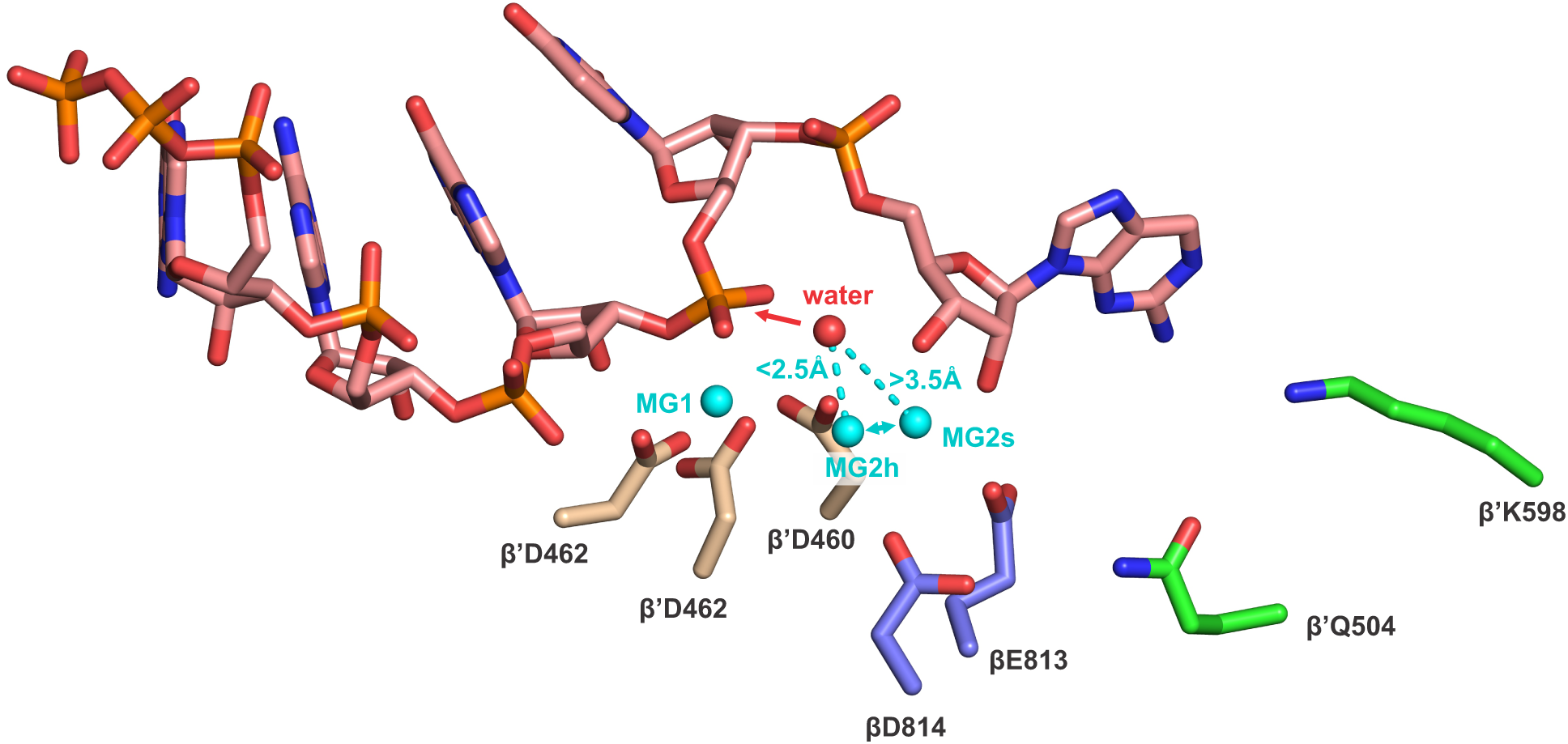
β’Q504R and β’K598W substitution may increase RNA cleavage activity by altering the position of MG2. The backtracked RNA, the side chains of the acidic residues implicated in Mg^2+^ binding and the β’Gln^504^ and βLys^598^ are shown as sticks and colored as in the main text **Figure 5B**. The high-affinity MG1 is present in majority of RNAP structures and is universally coordinated by the Asp triad (β’Asp^460^, β’Asp^462^ and β’Asp^464^). In contrast, MG2 has been observed in two distinct locations: in-between βGlu^813^ and β’Asp^460^ (here named MG2s for structural) or in-between βAsp^814^, β’Asp^460^ and β’Asp^462^ (here named MG2h for hydrolytic, see below). The two positions are 2-3 Å apart, are considered mutually exclusive, but can likely reversibly interconvert (i.e. are in equilibrium symbolized by the two-sided cyan arrow). The former binding mode has been mainly observed in bacterial RNAPs, whereas the latter in the yeast Pol II and ppGpp-bound bacterial RNAP. Both positions of MG2 have been suggested to play a catalytic role during nucleotide incorporation (1,2) and play a structural role in the tagetitoxin-and ppGpp-bound structures (74,82). However, only MG2h can conceivably activate a water molecule for the attack (symbolized by a red arrow) on the RNA phosphate, thereby catalyzing the RNA cleavage reaction (81). We suggest that MG2 predominantly occupies MG2s site in the wild-type 2AP-TEC leading to a stable 1-nt backtracked state with a low RNA cleavage activity. Arg introduced in position β’504 can form a salt bridge with βGlu^813^ and bias the equilibrium between the MG2s and MG2h towards the latter site thereby destabilizing the backtracked state, but increasing its RNA cleavage activity. β’K598W substitution may also increase the RNA cleavage activity by increasing the occupancy of the MG2h site. It may possibly do so by altering the overall pose (and thereby positions of the hydroxyl groups) of the backtracked nucleotide via contacts with its nucleobase. Figure was prepared using PyMOL Molecular Graphics System, Version 1.8.6.0 Schrodinger, LLC and atomic coordinates of the model of 1-nt backtracked state (the main text **Figure 5A,B**) supplemented with MG2h and the catalytic water molecule.

**Supplementary Figure S7.**
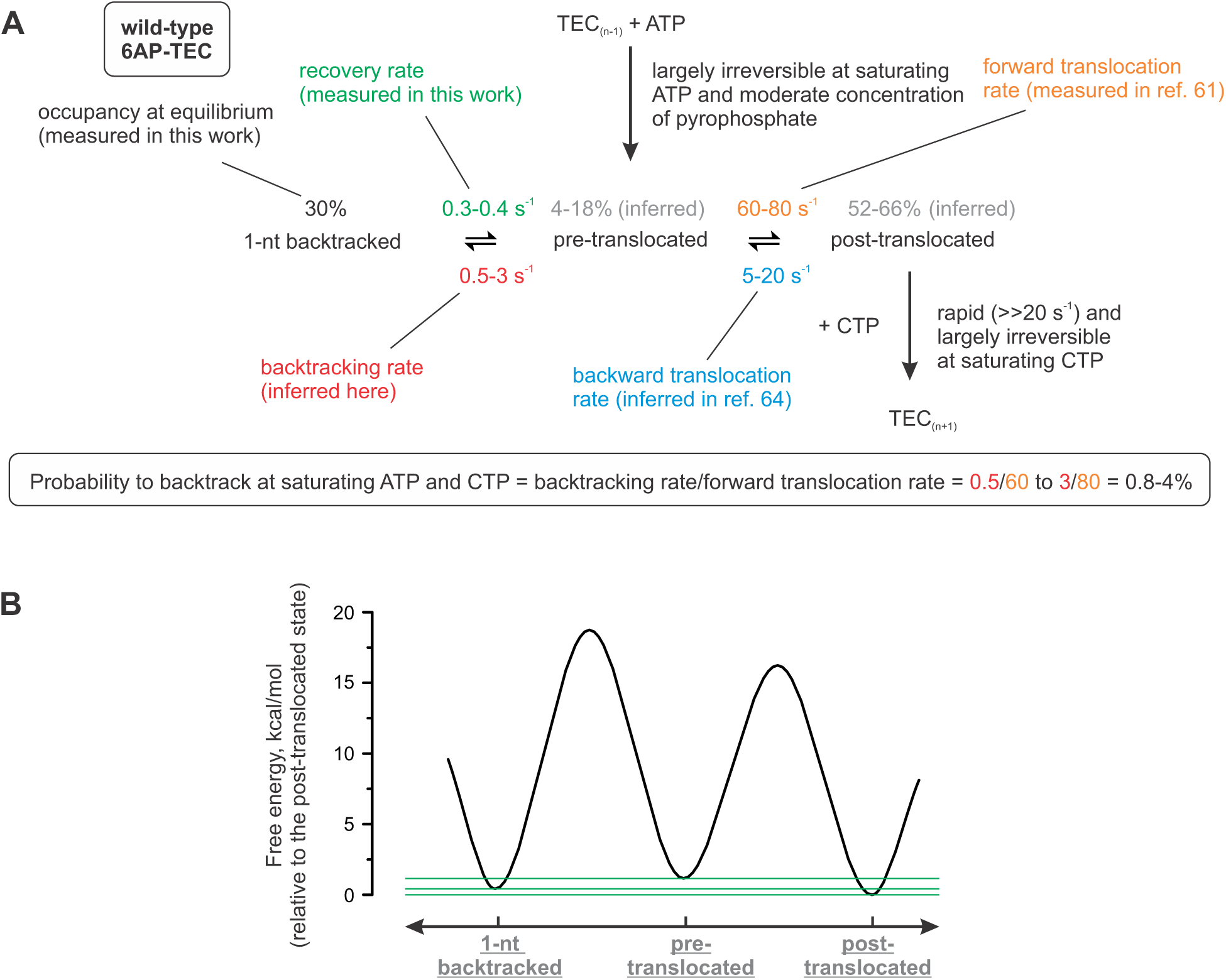
The backtracking probability during processive transcription through the sequence position corresponding to the 6AP-TEC. Simulations were performed using KinTek Explorer software (KinTek Corporation, Austin, TX). **(A)** We defined the backtracking probability at saturating concentrations of CTP and ATP as the ratio of the backtracking rate and the forward translocation rate. Backtracking rate (1-3 s^-1^) was inferred based on the other rates in the scheme and the equilibrium occupancy of the backtracked state. The equilibrium occupancy and the recovery rate of the backtracked state (28%, 0.3 s^-1^) were measured in this work. The forward translocation rate, (60-80 s^-1^) was measured by Malinen et al. 2012 (61). The backward translocation rate (5-20 s^-1^) was set based on the data reported by Malinen et al. 2014 (64). Specifically, we made three assumptions: (*i*) the wild-type, β’F773V and β’P750L RNAPs have similar backward translocation rates (stabilized TH slow forward, but are not expected to affect the backward translocation rate);(*ii*) the backward translocation rate is similar in the TECs with GMP and AMP at the RNA 3’ end (such TECs display similar fractions of the pre-translocated state; Figure 2A in ref. 64) and (*iii*) the forward and the backward translocation rates are similar at 1 mM and 10 mM Mg^2+^ (post-and pre-translocated states possess only high-affinity Mg^2+^ bound to the Asp triad). Mg^2+^ anchors the RNA 3’ end in the post-translocated register (1,61) and the incomplete saturation of the high-affinity site may result in a higher backward translocation rate. To account for the possibly higher backward translocation rate at 1 mM Mg^2+^ we increased the upper bound twofold: from 5-10 s^-1^ estimate at 10 mM Mg^2+^ in Malinen et al. 2014 (64) to 5-20 s^-1^ interval that we used to simulate the system at 1 mM Mg^2+^ (i.e. we assumed that *K*_*D*_ < 1 mM for Mg^2+^ bound to the Asp triad). **(B)** The energy profile for the equilibria in (A). Green horizontal lines aid the comparison of the energy minima.

